# Logic of optimal collective migration in heterogeneous tissues

**DOI:** 10.64898/2026.03.19.712843

**Authors:** Uday Ram Gubbala, Diana Pinheiro, Edouard Hannezo

## Abstract

Collective cell migration is a critical process in embryogenesis and cancer invasion. Recent work has shown that uniform tissues can undergo sharp rheological transitions, with collective motion emerging above a critical cell motility. In vivo, however, migration typically involves multiple populations with distinct motile and adhesive properties, and how this heterogeneity shapes collective dynamics remains unclear. Here, using two different vertex model implementations, we show that migration of heterogeneous clusters through tissues is maximized at intermediate adhesion strength: too little and the cluster fragments, too much and cluster cell cohesion suppresses the rearrangements needed for forward motion. We test our model against recent and new data on zebrafish mesendoderm invasion, where graded Nodal signalling regulates both motility and adhesion differences. By mapping measured Nodal levels to mechanical parameters, the model not only reproduces migration outcomes across homogeneous and heterogeneous clusters, but also discriminates between alternative adhesion rules. Strikingly, the inferred parameters place the system near the predicted optimum, where adhesion is strong enough to maintain cohesion yet graded enough to allow selective coupling among heterogeneous neighbors. These results identify an optimal balance between cohesion and interfacial remodeling as a general principle coordinating collective invasion in heterogeneous tissues.

**Significance statement:** Cells often migrate collectively during embryonic development and cancer invasion, but tissues are rarely uniform and different cells differ both in their adhesion and activity. Using models of tissue mechanics, we show that collective invasion is maximized at an intermediate level of adhesion within the migrating cluster cells: too little and the cluster falls apart, too much and it cannot advance. We test this principle against experiments in zebrafish gastrulation, where a signaling gradient simultaneously controls both cell motility and adhesion. The model reproduces migration outcomes across a range of experiments and identifies the adhesion rule cells use to selectively stick to neighbors. These results reveal a simple mechanical logic for how heterogeneous cell collectives coordinate invasion.

## 1 Introduction

Collective cell migration plays a central role in embryogenesis, tissue regeneration, and cancer invasion [**1–5**]. A particularly well-studied organizational mode is leader–follower migration, in which a subset of cells at the front generates protrusive forces and interprets external signals to guide the movement of less motile followers [**6–9**]. While classical studies emphasized biochemical guidance by leader cells [**8, 10, 11**], recent work has revealed a richer picture: followers exhibit diverse, active behaviors [**9**] and generate forces that promote leader emergence and sustain long-range coordination [**12, 13**], pointing to mechano-chemical couplings between these distinct cell populations during collective invasion. In parallel, efforts have been made to understand how even in homogeneous tissues, small changes in mechanical properties (e.g. cell-cell adhesion) or active forces (e.g. cell motility) can drive rheological transitions and collective migration [**14, 15**]. Indeed, inspired by the physics of jamming and glass transitions, biological tissues have been shown to transition between solid-like and fluid-like states [**16–18**]. Such tissue fluidization can underpin the onset of collective migration, as demonstrated in vitro in tumor collectives [**19**] or airway epithelium [**20**], and minimal biophysical theories, such as Vertex models, have been instrumental in predicting the onset of these transitions in homogeneous tissues [**21–24**]. Yet their application to actively migrating and mechanically heterogeneous collectives remains limited [**25–28**]. This motivates our central question: how do mechanical heterogeneity and local tissue rheology determine the leader–follower organization and migration efficiency of invading cellular clusters?

To address this, we developed a modeling framework linking interfacial mechanics with active migration in heterogeneous epithelial clusters. We considered two key sources of heterogeneity, motivated by recent experimental findings. Firstly, we assumed that different invading cells exert different active migration forces, as observed, for example, leader-follower organization in zebrafish gastrulation movements [**29**] or fibroblast-led collective invasion in tumors [**30**]. Secondly, a further source of mechanical heterogeneity in cellular collectives is the regulation of interfacial forces between distinct subpopulations within the cluster and at the cluster boundary. Such regulation can arise through differential expression of adhesion molecules or through actomyosin tension at heterotypic interfaces [**26, 31–33**]. To obtain generic and robust predictions, we used two complementary model implementations that incorporate both ingredients: the Self-Propelled Voronoi (SPV) model, which applies motility forces at cell centres and reconstructs cell shapes through a Voronoi tessellation [**24**], and the Active Vertex Model (AVM), which applies forces at vertices and explicitly resolves individual cell boundaries [**34, 35**]. This allowed us to systematically investigate how heterotypic tension, cluster composition, and motility heterogeneities influence collective migration dynamics. We uncover general mechanical principles that enable optimal invasion of heterogeneous clusters through solid-like surroundings and test these predictions using published and new datasets from zebrafish gastrulation [**29**].

## 2 Results

### 2.1 Vertex model of migrating cluster invasion

We set out to understand the physical principles that govern cell cluster migration using a two-dimensional vertex model (implemented in two complementary variants: SPV and AVM), a framework commonly employed to describe the mechanics of confluent tissues [**24, 34–36**]. In this model, we implemented a heterogeneous tissue in which a cluster of cells with distinct motilities and interfacial interactions was embedded within a tissue of non-motile cells, with identity-specific adhesive properties (Fig. 1a). This configuration recapitulates experimental settings such as zebrafish mesendoderm clusters, where high Nodal signalling activity promotes invasion into the surrounding ectoderm [**29**], as well as mixed tumor spheroids in which contractility-dependent cell sorting influences collective invasion [**32**]. To impose an impenetrable boundary, a thin strip of edge cells at the left boundary is held fixed; these cells provide the rigid edge against which invasion is measured (see Methods for details).

**Fig. 1.**
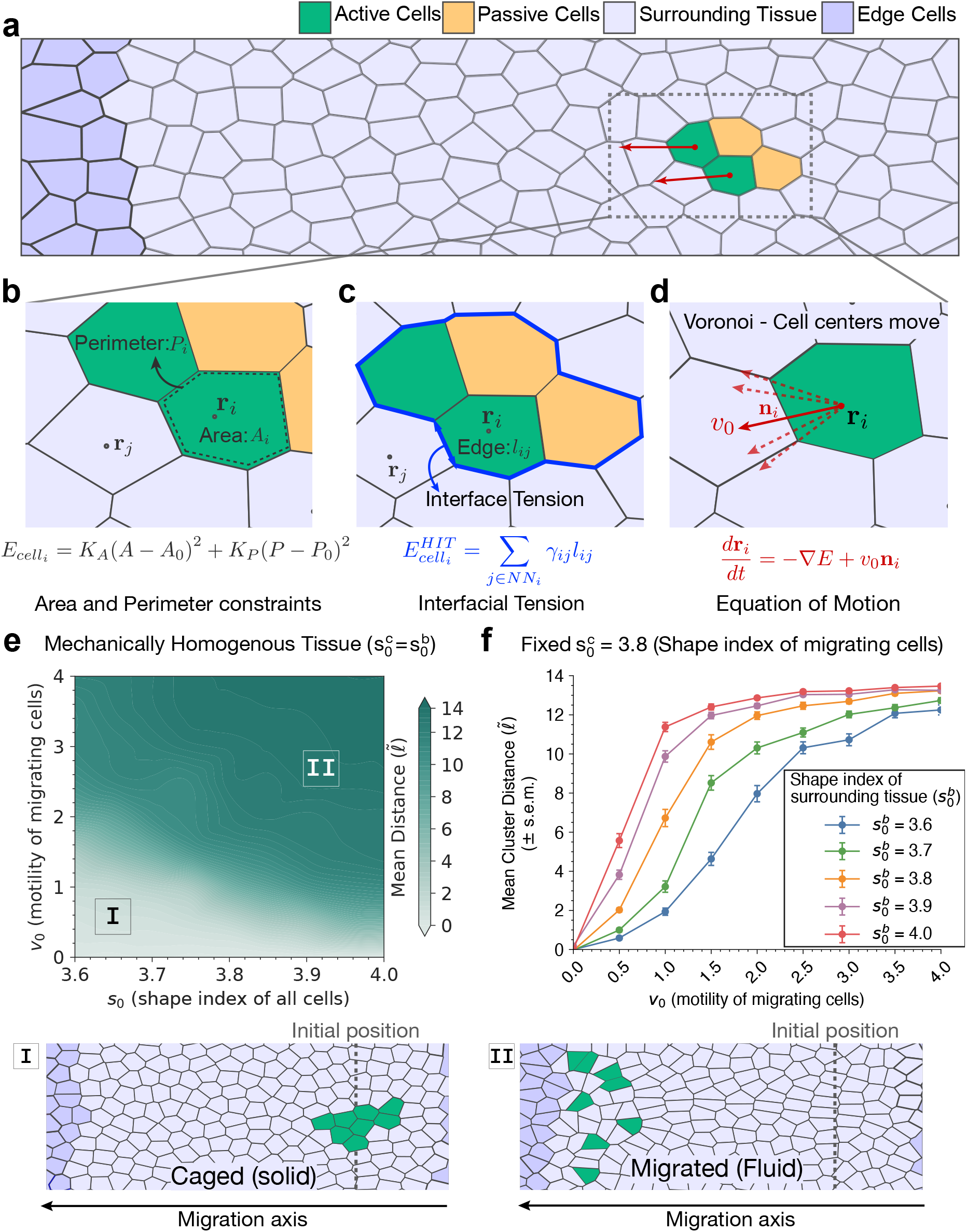
Vertex-based model and collective migration behavior. (a-d) Schematic representation of the different physical ingredients of the model. Representative confluent tissue (Voronoi tessellation, a) with a cluster of active and passive cells (resp. green and yellow) embedded in a surrounding tissue (purple). All cells evolve according to mechanical energy function (b), defined by area and perimeter constraints. We also consider different adhesion properties of different cell types (c), with for instance interfacial tension along cluster–tissue edges, which mediates effective cluster cohesion. Finally, cluster cells exert directed and noisy active motility forces (d), with overdamped dynamics. (e) Phase diagram of mean cluster migration distance in the (*v*_0_, *s*_0_) plane for homogeneous tissues 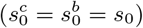. Migration increases with motility *v*_0_ and increasing *s*_0_. Final-time snapshots (I–II) show representative caged and migrated states. (f) Mean cluster distance versus motility *v*_0_ at fixed cluster shape index 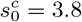 while varying the background shape index 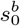 (legend). Larger 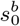 enables comparable displacement at lower *v*_0_.

In our vertex framework, each cell is represented as a polygon with area *A*_*i*_ and perimeter *P*_*i*_ (Fig. 1b), and the tissue mechanics are described by the energy

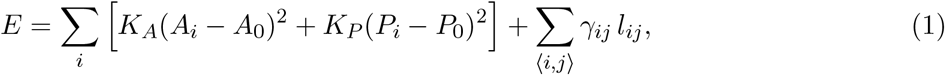

where *K*_*A*_ and *K*_*P*_ are the stiffnesses associated with area and perimeter deviations, and *γ*_*ij*_ is the line tension assigned to the edge of length *l*_*ij*_ shared by cells *i* and *j*. For the purpose of the interfacial mechanics, each cell is labeled by its tissue class *σ*_*i*_ ∈ {*c, b*}, where *c* denotes cells belonging to the migrating cluster and *b* denotes cells belonging to the surrounding background tissue. This cluster/background label is independent of any additional intra-cluster heterogeneity. We then define

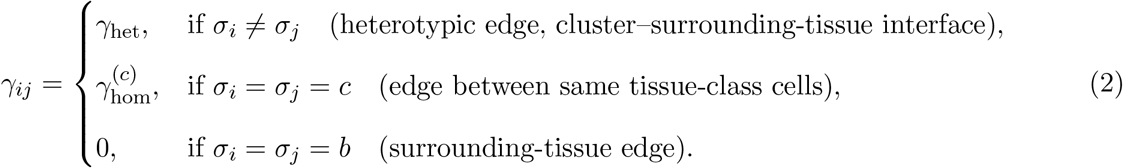

Unless otherwise stated, we set cluster–cluster and surrounding-tissue line tensions to zero (as spatially uniform shifts in line tension only renormalize the preferred cell perimeter; see the Supplementary Information (SI) for details) and vary only *γ*_het_. With this convention, *γ*_het_ measures the heterotypic line tension relative to the homotypic baseline. A positive *γ*_het_ penalizes heterotypic edge length and promotes contraction of cluster–surrounding-tissue interfaces, leading to sharper boundaries, whereas a negative *γ*_het_ makes heterotypic contacts energetically favorable relative to homotypic ones and therefore promotes interface expansion and cell mixing (Fig. 1c) [**26**]. Throughout, we refer to *γ* as an effective adhesion or interfacial-coupling parameter, while noting that in the vertex energy it enters explicitly as a relative line tension.

We characterize cell mechanics through the shape index 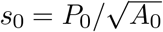, which captures the preferred cell geometry and has been shown to predict the rheological behavior of uniform tissues [**24, 37–39**]. The dynamics are overdamped and non-dimensionalized by setting *A*_0_ = 1, which fixes the length scale, and by measuring time in units of 1*/*(*µK*_*P*_ ), so that *µ* = 1 without loss of generality. Cluster and surrounding background tissue mechanics are varied through their respective shape indices 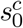 and 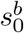, while motile cells are assigned an active motility of magnitude *v*_0_ along a polarity direction **n**_*i*_. This active drive is applied at cell centers in SPV (Fig. 1d) and at vertices in AVM (Supp. Fig. S1a), with the polarity angle evolving as an Ornstein–Uhlenbeck process of persistence time *τ* . Polarity vectors are independent across neighbouring cells and weakly biased along the −*x* direction, representing guided motion as observed in zebrafish mesendoderm and other chemotactic or mechanically guided migrations [**29, 40, 41**]; further model details are given in the Methods and SI.

As a starting point, we examined the SPV model (Fig. 1d). We first considered mechanically homogeneous tissues, keeping the cluster size fixed and assigning identical mechanical properties to both cluster and surrounding tissue 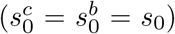. We focus on the mean cluster displacement at a fixed time as an operational measure of invasion, since it closely mirrors experimental readouts (see Methods for more information). The displacement map in the (*v*_0_, *s*_0_) parameter plane shows a gradual increase in invasion with motility strength *v*_0_ (Fig. 1e). In the fluid-like regime above the SPV rigidity transition [**24**] (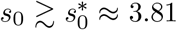), even modest *v*_0_ yields appreciable migration and the mean distance continues to rise with increasing activity. By contrast, below this transition 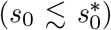, displacement remains small at low *v*_0_ and increases only once *v*_0_ becomes sufficiently large. This trend is consistent with motility-driven unjamming in confluent models [**24**]: increasing activity promotes rearrangements in the surrounding tissue and thereby enables sustained forward invasion.

We next introduced heterogeneity by varying the mechanical properties of the surrounding tissues while keeping migrating cluster mechanics fixed at 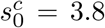. Increasing 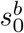 systematically enhanced invasion (Fig. 1f), with representative snapshots showing that clusters in fluid backgrounds travel farther and advance more cohesively. By contrast, varying the cluster shape index 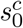 at constant background mechanics 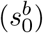 had little effect on invasion (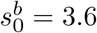 and 4.0 in Supp. Fig. S1d,e).

To test that these trends do not depend on a specific vertex-model implementation, we performed the same simulations in the Active Vertex Model (AVM) (Supp. Fig. S1a) and found the same qualitative results (Supp. Fig. S1b–g). A closer comparison of the displacement curves across both models (Fig. 1e,f and Supp. Fig. S1b–g) revealed quantitative differences between their dynamical responses: in SPV, invasion increases gradually with *v*_0_, whereas in AVM it rises more steeply and then plateaus, reflecting differences in degrees of freedom and rearrangement mechanics between the two models.

Despite these differences, the key result is robust across model implementations: invasion is primarily set by background mechanics, while the cluster’s shape index has limited predictive value in heterogeneous tissues, underscoring the need for caution when inferring mechanical properties from shape-based metrics as is commonly done in homogeneous settings [**42, 43**].

### 2.2 Optimal strength of cell-cell adhesion for cluster invasion

Motivated by our central question of how heterotypic interfacial forces control collective invasion, we introduced a heterotypic line tension *γ* along mixed cluster–host edges (Fig. 1c) and examined invasion efficiency for homogeneous bulk mechanics 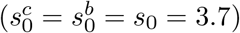. We chose this shape index to probe the effect of *γ* in a solid-like regime (*s*_0_ < 3.81), where invasion is limited by the ability of the surrounding tissue to undergo neighbour exchanges. We mapped mean cluster displacement across a parameter scan of *γ*, motility *v*_0_, and cluster size. Both SPV and AVM showed a peak in displacement at intermediate *γ*, and this optimal window became stronger and wider with increasing motility and initial cluster size (Fig. 2a,b). To illustrate the resulting migration regimes, we focus on *v*_0_ = 1.5 in SPV, for which cluster displacement at *γ* = 0 remains limited at *s*_0_ = 3.7 (Fig. 1e,f), and the corresponding *v*_0_ = 0.75 in AVM (Supp. Fig. S1b,c). In both implementations, increasing *γ* reveals three migration regimes with a clear optimum at intermediate *γ* (Fig. 2c,d), and these trends remain robust under parameter changes (Supp. Fig. S2c,d,g,h). The optimal *γ* window becomes increasingly pronounced with cluster size, consistent with a collective invasion advantage over single-cell motion (Fig. 2c,d; Supp. Fig. S3a,e). Together, these results identify an optimal range of heterotypic tension that maximizes invasion in solid-like surroundings.

**Fig. 2.**
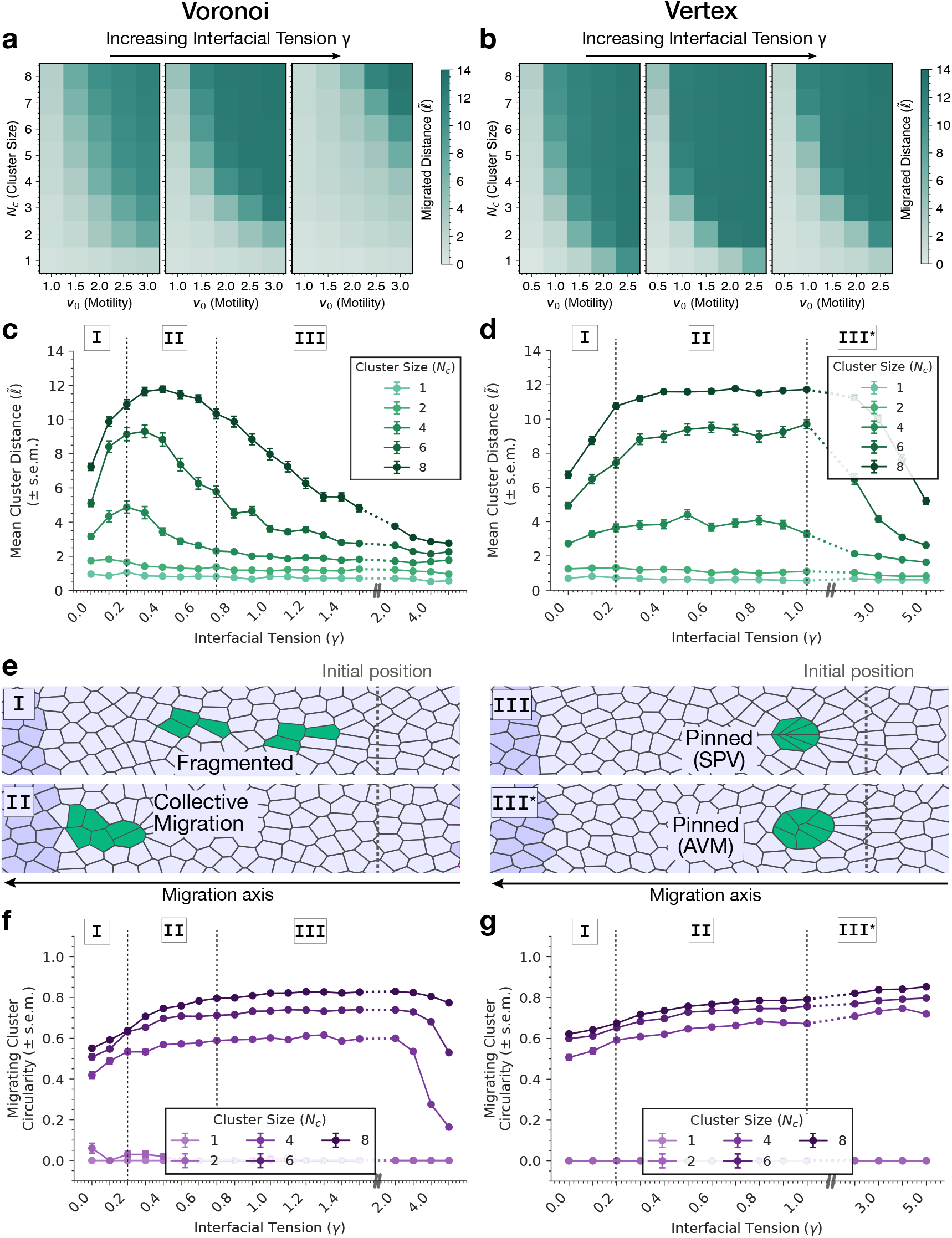
Optimal heterotypic tension for collective invasion. (a,b) Mean cluster migration distance as a function of migration forces *v*_0_ and initial cluster size *N*_*c*_, for the SPV/Voronoi (a) and AVM/Vertex (b) models, for increasing heterotypic interfacial tension *γ* (from left to right panels, *γ* = 0, 0.3, 5 for SPV; *γ* = 0, 0.5, 5 for AVM). (c,d) Mean cluster distance versus heterotypic tension *γ* at fixed motility (*v*_0_ = 1.5 for SPV and *v*_0_ = 0.75 for AVM) for different initial cluster sizes *N*_*c*_. In both models, invasion depends non-monotonically on *γ* and is maximized at intermediate values of adhesion. Dashed vertical lines indicate representative *γ* values for the distinct invasion regimes shown in panel (e). (e) Representative final configurations illustrating the regimes identified in (c,d): fragmented clusters are observed at low *γ* (I), while cohesive collective migration occurs at intermediate *γ* (II), both for AVM and SPV models (panels here show SPV). For high *γ* pinned states, migration is impaired in both models, although with different geometries: a compact pinned cluster in SPV (III) and a rounded pinned cluster in AVM (III^*\**^). (f,g) Cluster circularity versus heterotypic tension *γ* under the same parameter regime as in (c,d) for the SPV (f) and AVM (g) models. In SPV, circularity rises at intermediate *γ* and decreases again at high *γ*, whereas in AVM it continues to increase, reflecting distinct high-*γ* cluster morphologies in the two models.

We next asked what is the physical basis of the optimal *γ* for migration. Snapshots from both models (Fig. 2e) revealed three regimes: at low *γ*, the cluster–background interface is highly deformable and cluster cohesion is weak; at intermediate *γ*, the interface is stabilised and clusters move more efficiently as a compact group; in contrast, at high *γ*, clusters become rounded and kinetically ‘pinned’ (i.e., they advance only weakly despite sustained motility; in contrast to *caging*, where motion is arrested because a solid-like background suppresses rearrangements in the bulk).

To quantify these changes in cluster organization, we computed several cluster morphometrics, including cluster circularity (Fig. 2f,g), the weighted cluster size (the mean size of connected components weighted by their contribution to the total cluster population at the final time point; Supp. Fig. S2a,e), and the associated changes in average perimeter and area (Supp. Fig. S4a–h). In SPV (Fig. 2f), circularity rises sharply from *γ* = 0 until it reaches a plateau, and decays only at very large *γ*, where strong pinning and highly compact shapes emerge, consistent with the sharp reduction in cluster-cell perimeter and area (Supp. Fig. S4a–d). In AVM, circularity gradually increases with *γ* and remains high even for large *γ* (Fig. 2g), while perimeter and area decrease more gradually over the same range (Supp. Fig. S4e–h). Weighted cluster size measurements (Supp. Fig. S2a,e) show the same qualitative trend in both models: they are lowest at *γ* = 0, where extensive dispersion occurs, increase rapidly as the interface is stabilised, and then saturate once cohesion is established. Together, these morphology-based metrics indicate that increasing *γ* from 0 to moderate values primarily suppresses dispersion and promotes cohesive, rounded clusters, whereas the loss of invasion at large *γ* occurs even though circularity and effective cluster size remain high.

Interestingly, the suppression of cluster dispersion occurred once the interfacial tension reached a threshold of *γ* ≃ 0.3–0.4 (for *N*_*c*_ = 8), which coincided with the rise and peak of the mean cluster displacement (Fig. 2c,d) and with the increase in weighted cluster size (Supp. Fig. S2a,e). We therefore hypothesized that the beneficial effect of *γ* at small tension values is a collective effect mediated by the fact that effective cluster size remains larger as adhesion increases. In line with this, displacement in single-cell clusters was small and did not show any increase with *γ* (Supp. Fig. S3a,e; see SI for more details), whereas clusters systematically migrated farther with increasing initial size (Fig. 2c,d). To test this quantitatively, we plotted weighted cluster size versus displacement distance across initial cluster sizes and interfacial tensions, which revealed that displacement increases with effective cluster size (Supp. Fig. S2b,f). This indicates that a first key contribution of small *γ* is to maintain cohesive clusters, allowing cells to coordinate their motility and collectively exert a larger net driving force on the surrounding tissue.

To quantify this effect further, we computed front-edge creation and rear-edge detachment rates, which measure how often new heterotypic contacts form at the leading edge of the cluster and how often existing contacts are lost at the trailing side (see Methods for precise definitions). These events are realised through local four-cell T1 transitions [**21, 36**] at the cluster interface. At *γ* = 0, mixed interfaces rearrange freely because there is no energy penalty for rearrangements (Eq. 1): homotypic contacts break easily and the rates of front-edge creation and rear-edge loss remain low (Supp. Fig. S3c,d,g,h), consistent with a shallow barrier to interfacial rearrangements. For small but non-zero values of *γ*, the relative energy cost of homotypic rearrangements increases, so the creation and loss rates of heterotypic edges at the front and rear rise and peak at the same intermediate *γ* that maximises migration (Supp. Fig. S3c,d,g,h). In this regime, motility-generated cluster polarity can be converted into sustained forward motion because the interface is stable enough to hold the cluster together yet still flexible enough to easily remodel and undergo directed rearrangements. However, when *γ* becomes too large, this balance reverses. Strong *γ* favours short heterotypic edges and promotes the formation of multifold vertices at the cluster interface, a behaviour reported in both Voronoi and vertex-based descriptions [**26, 33**]. In a migrating cluster, neighbouring heterotypic edges then pull against one another in a geometric tug-of-war in order to shorten, strongly suppressing the T1 transitions required for forward motion and driving both rear-edge detachment and front-edge creation sharply down (Supp. Fig. S3c,d,g,h). Although rearrangements in the AVM are triggered through an explicit edge threshold, the same qualitative picture appears: edge creation and loss remain possible at intermediate *γ* but are strongly reduced once line tension dominates. Altogether, this explains why intermediate values of *γ* allow for optimal migration, by balancing propulsive forces with an interface that is cohesive yet still able to rearrange.

### 2.3 Optimal adhesive coupling for leader–follower transport

Next, we sought to model leader–follower dynamics as a third source of heterogeneity in the system (Fig. 1a,b). In particular, we considered clusters composed of actively migrating leaders and passive followers with *v*_0_ = 0, to ask how motile cells transport non-motile ones. This setup is motivated by zebrafish gastrulation [**29**], where highly protrusive leaders pull on less migratory followers, as well as by previous reports showing mechanical interactions between cancer-associated fibroblasts and tumour cells [**30**]. Since SPV and AVM yielded similar invasion phenomenology in the previous section, we focus here on the SPV formulation for simplicity. We assigned *γ*_1_ to leader–tissue edges and *γ*_2_ to follower–tissue edges while keeping leader–follower edges neutral, and focused on symmetric coupling by setting *γ*_1_ = *γ*_2_ ≡ *γ*, i.e. leaders and followers experience the same interfacial coupling to the surrounding tissue (Fig. 1b).

We quantified leader–follower transport using two complementary metrics: the distance travelled by passive cells and the leader–follower displacement gap (see Methods for definitions). For equal-proportion leader–follower clusters, follower displacement depended non-monotonically on *γ*, with a clear optimum at intermediate coupling (Fig. 3a). At low *γ*, leaders move ahead without effectively carrying followers, leading to an uncoordinated regime in which leaders and followers separate and follower transport is weak (Fig. 3a–c, I). At intermediate *γ*, interfacial coupling is sufficient to transmit leader-driven motion to followers, producing optimal transport in which active and passive cells co-migrate as a cohesive unit, with maximal follower displacement and improved coordination (Fig. 3a–c, II). For large *γ*, migration is strongly suppressed as the cluster–background interface becomes pinned, reducing follower displacement even though leaders and followers remain close (Fig. 3a–c, III). Cluster composition also matters: at fixed total initial cluster size, increasing the number of active cells boosts follower displacement and improves leader–follower coordination (Supp. Fig. S5a,b), indicating that the effective driving of the cluster is collectively set by the total number of motile cells.

**Fig. 3.**
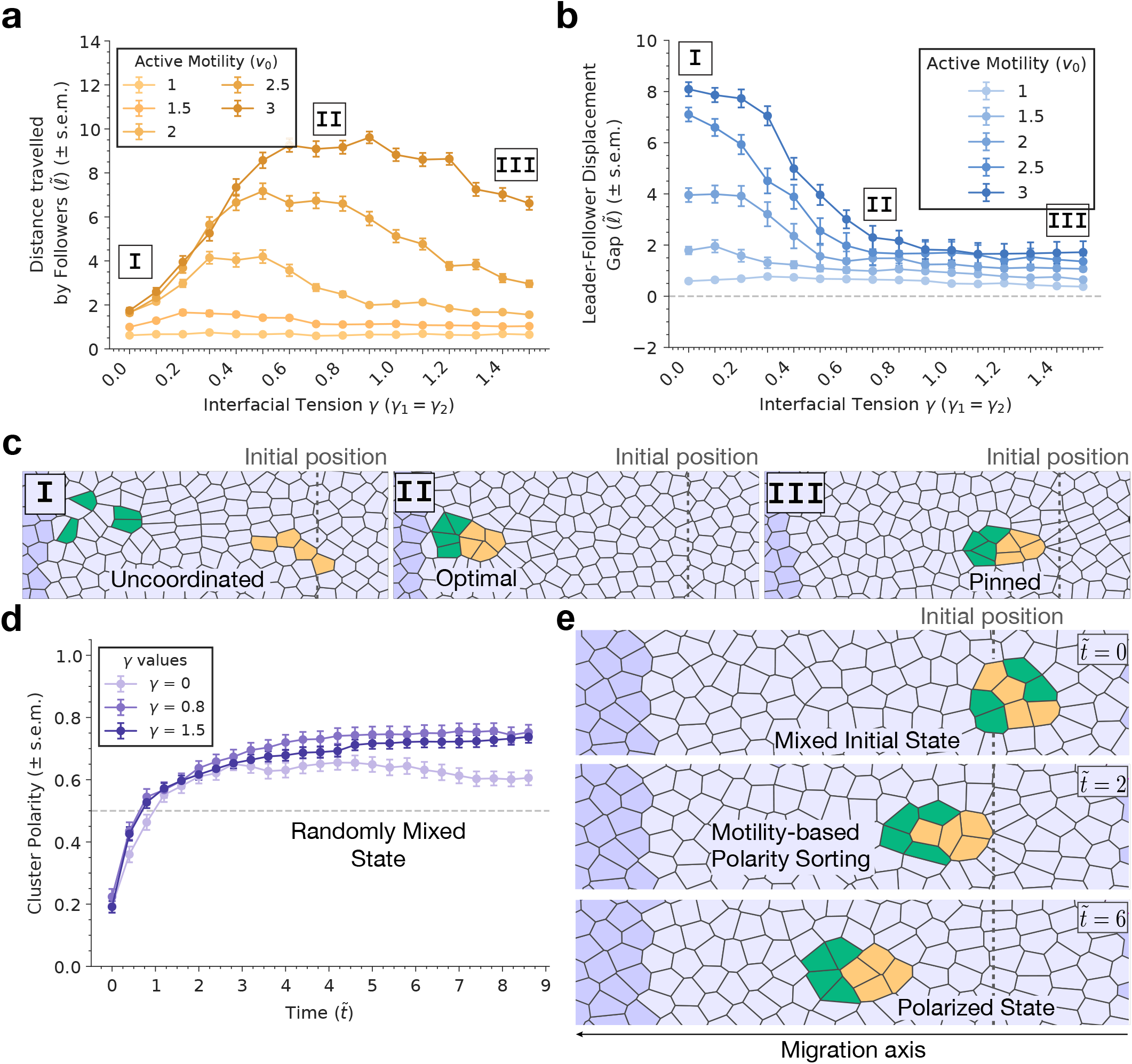
Leader–follower transport and polarity sorting in active heterogeneous clusters. (a-c) Simulation outputs for heterogeneous clusters (leaders:followers ratio = 1 : 1) in a solid background tissue (fixed cell shape index *s*_0_ = 3.7). The mean distance travelled by follower cells increases non-monotonously as a function of interfacial tension *γ*, as shown in (a) different active motilities *v*_0_ (legend). This occurs due to leader-follower detachments at low *γ*, as quantified by the leader–follower gap versus interfacial tension *γ* (b). We show representative cluster configurations (c) corresponding to the three regimes identified in (a,b): uncoordinated leader-follower motion at low *γ* (I), optimal follower transport at intermediate *γ* (II), and pinned states at high *γ* (III). (d) Cluster polarity, defined from the relative positions of leaders and followers along the cluster front–rear axis, as a function of time for three values of *γ*, showing robust sorting. (e) Time sequence illustrating motility-based polarity sorting from a mixed initial state to a polarized configuration.

Heterogeneities in cell motility also lead to self-organization within mixed clusters: starting from an initially mixed state, motile leaders progressively accumulate at the front, generating a polarized configuration with leaders enriched at the leading edge (Fig. 3d,e). This polarity build-up was observed across interfacial coupling strengths (Fig. 3d) and remained robust across motility and composition choices (Supp. Fig. S5c,d). Mechanistically, this robustness reflects the fact that line tension acts only on cluster–background edges: even when the outer interface is pinned, leader–follower contacts inside the cluster remain neutral, allowing motile cells to rearrange relative to followers and accumulate at the front.

### 2.4 Interplay between active migration forces and Nodal-dependent cell interactions in zebrafish gastrulation

We next asked whether the main features of our model (Fig. 4a) can capture mesendoderm migration during zebrafish gastrulation (Supp. Fig. S7a). Previous work showed that both motility and adhesion are regulated by a Nodal signalling gradient [**29, 44**], so that different mesendoderm progenitors with distinct Nodal signalling levels *N* have different mechanical properties. This was established via transplantation experiments of single or multiple migrating populations into Nodal-deficient host tissues to assess their invasion capabilities, providing a minimal assay to understand the dynamics of heterogeneous cluster invasion *in vivo*.

**Fig. 4.**
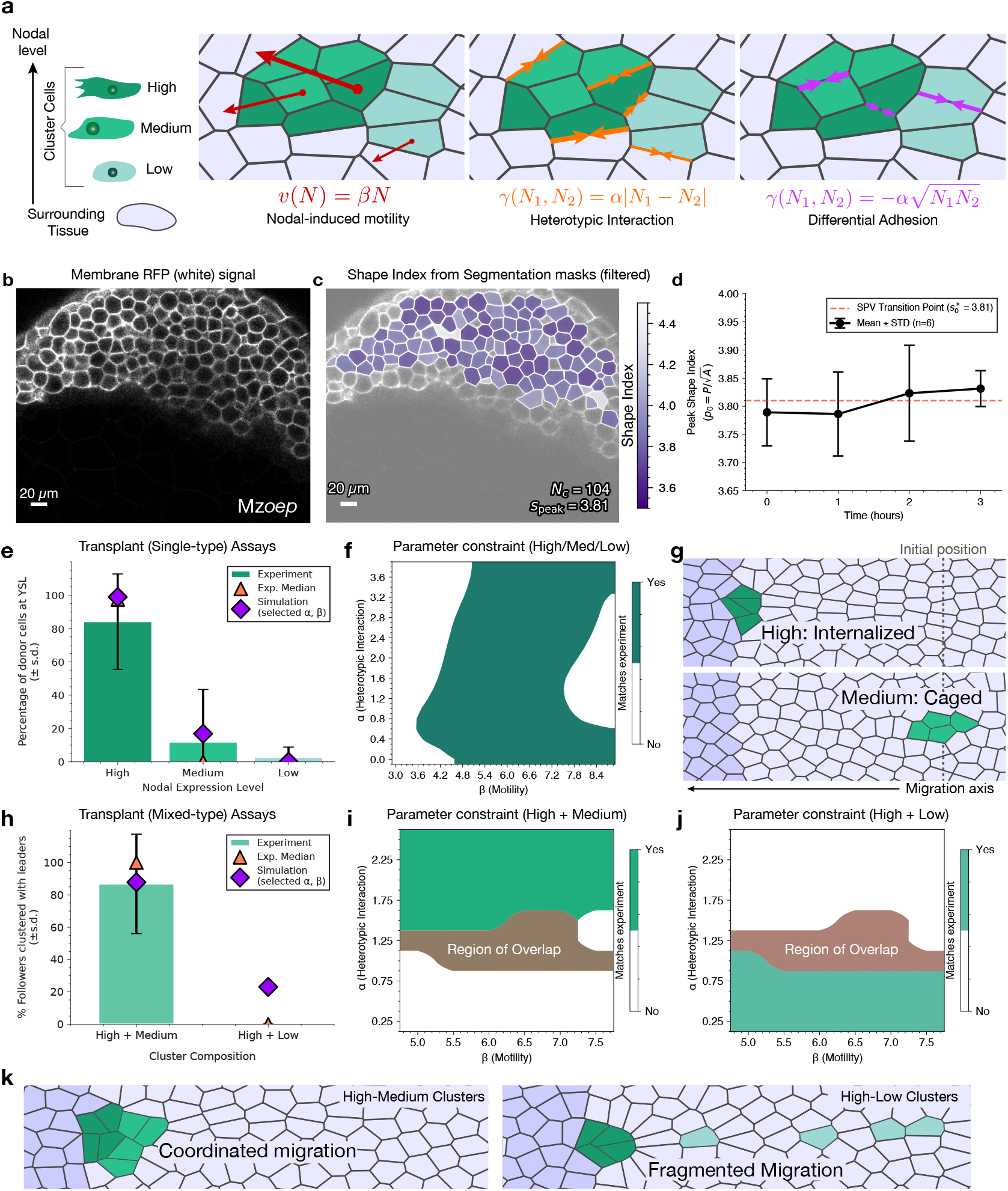
Testing the model against zebrafish mesendoderm invasion experiments. (a) Schematic of the model assumptions. Nodal level is mapped to cell motility through *v*(*N* ) = *βN*, and two candidate Nodal-dependent line-tension rules are considered: a heterotypic rule *γ*(*N*_1_, *N*_2_) = *α*|*N*_1_ *− N*_2_| and a differential rule 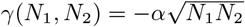. (b-d) Experimental estimation of the tissue shape index from segmentation masks. Panel (b) shows an example membrane signal, panel (c) the corresponding analyzed shape-index map, and panel (d) the time course of the measured peak shape index, which remains close to *s*_peak_ *≈* 3.81 over the observed time window. (e-g) Migration capability of clusters with different levels of Nodal signalling. In experiments (e), Nodal-High clusters (from 50% epiboly embryos) reach the YSL efficiently, whereas Nodal-Medium (from shield-stage embryos) and Nodal-Low (from 75% epiboly embryos) clusters rarely internalize [**29**]. This behaviour is captured by a finite region of (*α, β*) parameter space under the heterotypic rule (f). Representative simulations at the selected parameter values *α* = 1.25 and *β* = 6.5 are shown in (g). (h-k) Migration behavior of heterogeneous clusters with mixed Nodal levels. In experiments (h), the fraction of follower cells that remain clustered with leaders at the YSL is high for High+Medium donors but low for High+Low donors. The corresponding parameter constraints are shown in panels (i,j); their overlap identifies parameter values consistent with both mixed-cluster assays. Representative simulations at *α* = 1.25 and *β* = 6.5 are shown in (k), yielding coordinated migration for High–Medium clusters and fragmented migration for High–Low clusters. Scale bars: 20 *µ*m in (b,c).

Given our theoretical findings on the importance of surrounding tissue rheology for invasion, we first inferred its mechanical state by automatically segmenting cell shape of Nodal-deficient tissues from 2D sections (details in the Supplementary Information). The measured shape-index (peak), defined as 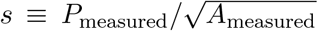 (as distinct from the preferred shape index *s*_0_ entering the vertex-model energy), was consistently around *s* ≈ 3.81, consistent with a jammed, solid-like tissue (Fig. 4b–d). We therefore modeled the surrounding tissue in the solid regime by setting *s*_0_ = 3.7, which yields observed cell shapes *s* close to the measured value [**24**]. In this regime, invasion requires cluster motility to overcome the mechanical resistance of the surrounding tissue and trigger local unjamming. This is consistent with previous work [**29**], which showed that only Nodal-high clusters (i.e. leaders) can invade, while Nodal-medium and -low clusters remain jammed (i.e. followers). To parametrize this, we assumed, as seen in experiments, that cluster active motility scaled linearly with Nodal levels as *v*_0_ = *βN* (Fig. 4a), and extracted Nodal signalling levels directly from experiments (which consider three populations: *N*_High_ = 0.44, *N*_Medium_ = 0.20 and *N*_Low_ = 0.08, Fig. S8a).

To test the model prediction that invasion relies not only on cluster active forces, but also the mechanical state of the surrounding tissue, we performed new transplantation experiments using host tissues with lower contractility (CA-Mypt, see Methods and SI for more information). This perturbation was previously shown to fluidize tissues [**45**] and we checked that it indeed impacts morphogenesis (Supp. Fig. S7b,c). Strikingly, when transplanting wild-type cells within this fluidized environment, we found that not only Nodal-high cells, but also Nodal-medium cells, could invade (Supp. Fig. S7e,f), consistent with the model (Fig. 1e,f). This suggests that zebrafish gastrulation is an ideal system to test how mechanical coupling within heterogeneous clusters shapes their ability to overcome local jamming.

We then turned to understanding the adhesive rules of heterogeneous clusters. Interestingly, Nodal has recently been reported to have multiple functions related to adhesion. On one hand, Nodal signalling can promote cell–cell contact stability and adhesion strength in early zebrafish tissues [**44, 45**]. This can be incorporated in the model via a differential adhesion rule, in which each cell *i* has an intrinsic adhesion strength proportional to signalling levels *N*_*i*_, so that adhesion at contacts between cells *i* and *j* is averaged, 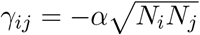 (Fig. 4a) [**46**]. However, adhesion between cells has also been proposed to depend on differences, rather than absolute levels, of Nodal signalling [**29**]. Mathematically, this corresponds to a heterotypic interaction rule, which is context-dependent and assigns an interfacial coupling between two cells *i* and *j* according to the difference in their Nodal levels, *γ*_*ij*_ = *α*|*N*_*i*_ − *N*_*j*_| (Fig. 4a). Under this rule, cells with similar Nodal levels remain more strongly coupled, whereas interfaces between cells with more distinct signalling states are associated with larger interfacial tension. This type of interaction has been suggested to be highly efficient at driving phase separation in immobile binary cell populations [**31**].

Given this multifaceted role of Nodal in setting up adhesion properties, we set out to quantitatively compare the design principles of both rules for efficient migration of heterogeneous clusters. Note that in either case, the entire mechanics of invasion depends only on two free parameters, *α* and *β*, dictating adhesion and motility strengths respectively.

We first examined simulations with a heterotypic interaction rule. The single-type transplantation assays already define a substantial admissible region in (*α, β*) space, since they require Nodal-high clusters to invade efficiently whereas Nodal-medium and Nodal-low clusters remain largely non-migratory (Fig. 4e–g; full parameter maps in Supp. Fig. S9a–f). As expected from the invasion optimum identified in Fig. 2, these assays constrain not only the motility scale *β*, but also the adhesion scale *α*: if *α* is too small, donor clusters lose cohesion and fragment, whereas if *α* is too large, strong donor– host coupling suppresses efficient invasion. The homogeneous assays therefore already select a broad intermediate range of *α*, while constraining *β* more sharply through the requirement that only the Nodal-high population invades.

Interestingly, Ref. [**29**] also performed migration assays combining populations with different Nodal levels, allowing us to test our predictions for heterogeneous clusters more directly. In particular, mixed clusters of Nodal-high and Nodal-medium cells efficiently co-migrate, whereas mixtures of Nodal-high and Nodal-low cells break apart, with Nodal-low cells remaining caged (Fig. 4h). First, we reanalyzed the dynamics of Nodal-high and Nodal-medium cluster experiments [**29**], which showed that, although the initial relative positions of leaders and followers are random, the clusters display the same motility-driven sorting behavior predicted in Fig. 3d,e, with Nodal-high cells reorganizing toward the leading edge during invasion (Supp. Fig. S8g,h).

Second, we performed systematic joint fitting of the Nodal-high+ Nodal-medium as well as Nodal-high+Nodal-low cluster experiments. We found that this strongly constrained the adhesion strength *α*: low *α* predicted that all mixtures should break apart, with only Nodal-high leaders migrating, whereas high *α* caused followers (both Nodal-medium and Nodal-low cells) to cage or pin the leaders, preventing global migration (Fig. 4h–k; Supp. Fig. S10a–f). By fitting median experimental outcomes and simulations (see Supp. Fig. S11 and SI for the corresponding distributional analysis), we obtained a tight intermediate parameter regime that reproduced all observations.

At the selected parameter values *α* = 1.25 and *β* = 6.5, inserting the measured Nodal levels into the heterotypic rule *γ*_*ij*_ = *α*|*N*_*i*_ − *N*_*j*_| (Fig. S8a) yields *γ*_H-M_ ≈ 0.30, *γ*_M-L_ ≈ 0.15, and *γ*_H-L_ ≈ 0.45. These values place interfaces between nearby Nodal levels (High–Medium and Medium–Low) close to the intermediate adhesion regime identified theoretically for heterogeneous clusters (Fig. 3a–c). By contrast, contacts between cells with very different Nodal levels (High–Low contacts) are weaker, allowing for strong yet selective mechanical interactions between different populations.

To challenge the model further, we turned to triple-transplant assays containing Nodal High, Medium, and Low populations. Experimentally, inserting Medium cells between High and Low populations partially restores collective behaviour, allowing a substantial fraction of low-Nodal cells to remain clustered with the leaders during invasion (Fig. 5a; [**29**]). The model reproduces this rescue robustly under the heterotypic interaction rule: a finite region of parameter space remains consistent with the triple-transplant assay while also satisfying the constraints from the previous homogeneous and mixed-cluster experiments (Fig. 5b), and representative simulations in this regime show collective internalization of High, Medium, and Low cells (Fig. 5d; Supp. Fig. S12a–c). This behaviour arises naturally because contacts between cells with similar Nodal levels (High–Medium and Medium–Low) remain sufficiently coupled to maintain cluster-wide cooperativity.

**Fig. 5.**
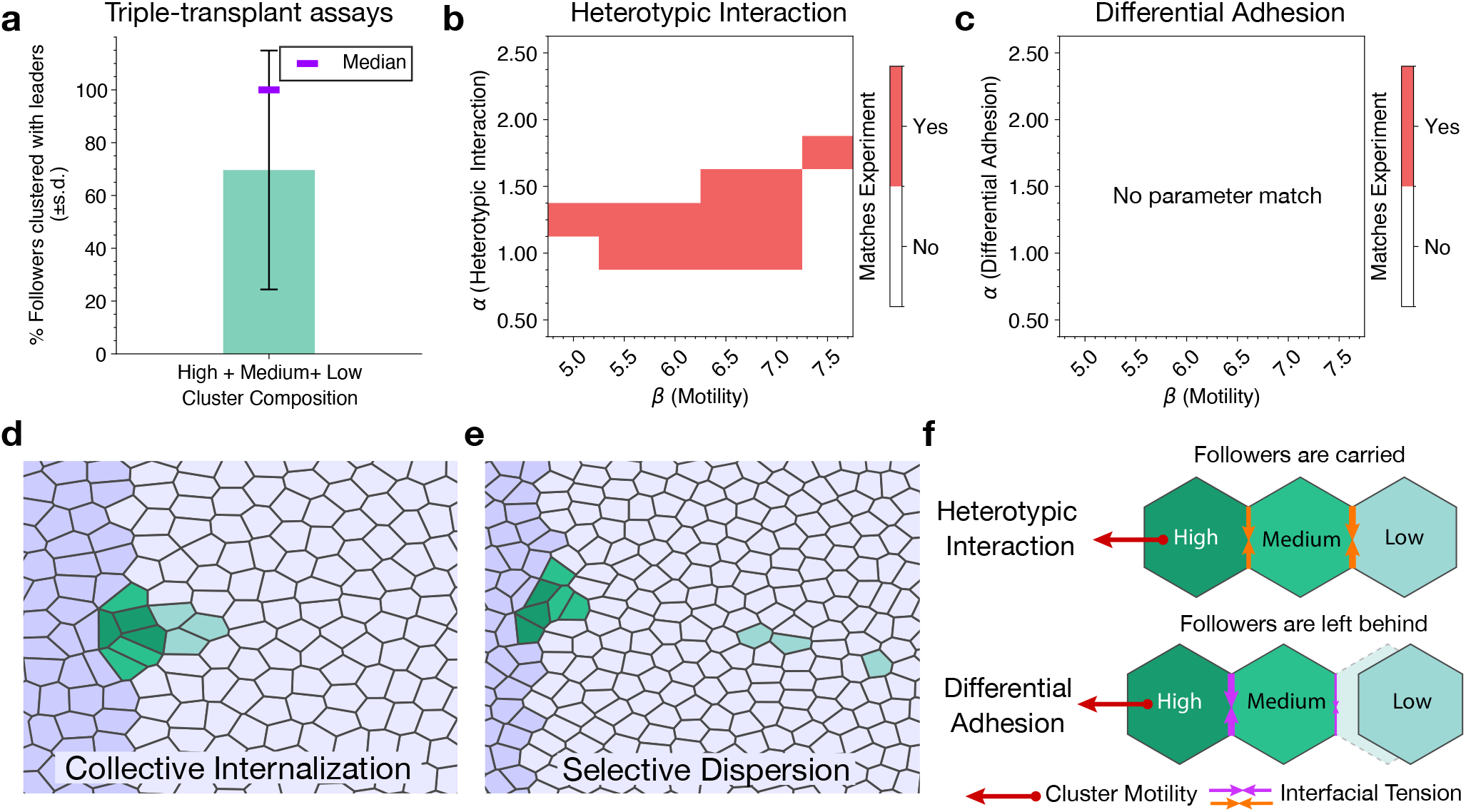
Three-cell-type cluster assays favor a heterotypic interaction rule. (a) Triple-transplant experiments with High, Medium, and Low Nodal cells. A substantial fraction of late cells remains clustered with leaders, indicating partial rescue of collective behaviour by the intermediate Nodal population. (b,c) Parameter pairs (*α, β*) that reproduce the triple-transplant assay under the two candidate interaction rules. Under the heterotypic interaction rule (b), a finite parameter region remains consistent with the experimental triple-transplant outcome and also lies within the parameter regime selected by the previous homogeneous and mixed-cluster assays. By contrast, under the differential-adhesion rule (c), no parameter pair satisfies the full set of constraints. (d,e) Representative simulations at *α* = 1.25 and *β* = 6.5. Under the heterotypic interaction rule (d), High, Medium, and Low cells undergo collective internalization, whereas under the differential-adhesion rule (e), the cluster disperses selectively. (f) Schematic summary contrasting the two rules. Heterotypic interactions promote coupling across unlike neighbors, allowing follower populations to be carried together during migration, whereas differential adhesion favors preferential cohesion among similar cell types, leading to selective dispersion and explaining why no single parameter set reproduces all experimental constraints.

Finally, we compared these findings to simulations with the differential adhesion rule, where adhesion strength is cell-intrinsic. We performed the same fitting procedure, first concentrating on the homogeneous and two-type clusters. We found that these datasets could still be reproduced by the model, although within a narrower parameter range (Fig. S8c,e,f; Supp. Figs. S9g–l and S10g–l; see SI text for details). Yet, under the differential adhesion rule, we could not find a robust overlap region that supports collective motion of triple transplants while also matching the homogeneous and mixed-cluster behaviors (Fig. 5c,f,h; Supp. Fig. S12d–f). This is because Medium–Low adhesion is always, by definition, weaker than High–Low adhesion, which prevents the formation of a stable adhesive chain from leaders to followers. Overall, these analyses support a Nodal-dependent heterotypic interaction rule that generates graded yet robust coupling between cells of adjacent Nodal levels, maintaining collective coherence within the cluster despite heterogeneity in both signalling activity and mechanical properties.

## 3 Discussion

Our results indicate that collective invasion in heterogeneous tissues is most effective when heterotypic adhesion lies in an intermediate range. At this level, cohesion is sufficient to hold the cluster together, allowing for a coordination of active forces, yet not so strong that neighbour exchanges are suppressed. This balance between cohesion and flexibility provides a simple physical rationale for efficient collective migration.

Comparing two complementary mechanical descriptions, the Self Propelled Voronoi (SPV) and the Active Vertex Model (AVM), we find consistent behaviour despite their different degrees of freedom, with both producing maximal invasion at intermediate adhesion. The SPV shows a sharper rise and a pronounced loss of motion at large interfacial tension, while the AVM maintains effective invasion across a broader parameter range. These quantitative differences reflect the implementation of interface remodelling and the accessible degrees of freedom in each formalism. At large interfacial tension, the arrest we observe in both models is reminiscent of pinning phenomena in driven interfaces, where increased line energy or local heterogeneity suppresses advance until a depinning threshold is crossed [**47**].

Anchoring the model to data on zebrafish gastrulation, we used imaging and cell segmentation to verify that the surrounding tissue is in a solid regime, which predicts that a critical motility is required for invasion, consistent with data. Mapping Nodal signalling levels to motility and interfacial tension, the model reproduced the experimentally observed outcomes: Nodal High clusters invade, whereas Nodal Medium and Nodal Low clusters do not (Fig. 4). Simulations of mixed clusters further matched experimental behaviour: Nodal High–Medium clusters are close to optimal cohesion, and therefore migrated with a leader–follower organization, while Nodal High–Low groups tended to separate, with the leaders advancing and the followers remaining jammed (Fig. 4; Supp. Fig. S8). The optimal adhesion regime that we uncover therefore helps migrating cells to robustly distinguish their followers, ensuring a combination of tight yet selective adhesion rules.

We expect these principles to extend beyond zebrafish gastrulation. Related leader–follower dynamics are reported when cancer associated fibroblasts pull tumor cells into stroma [**30, 48, 49**], during collective boundary crossing such as Drosophila border cell migration and vertebrate neural crest streams [**50–52**], or in heterogeneous organoid systems where specialized leaders guide group motion [**53, 54**]. In each of these contexts, regulated interaction and motility differences are likely to be key drivers shaping collective behaviour, suggesting that balancing cohesion and deformability may be a general principle for efficient invasion.

In the future, the model could be extended to include other interactions or mechanistic details. For instance, we modelled polarity as an Ornstein-Uhlenbeck process without feedback from shape or interface mechanics, based on experimental observations in zebrafish gastrulation that even followers are strongly polarized [**29**]. Yet, such feedbacks can be important in other epithelial collectives and migrating tissues [**55–57**]. Differences between SPV and AVM at large interfacial tension also point to model specific remodelling rules, which will benefit from closer study at higher resolution considering the detailed mechanics of the cytoskeleton and adhesive machinery at cell-cell junctions. Finally, our analysis focused on two-dimensional confluent sheets, although extending the framework to curved and three-dimensional settings could extend the applicability of the model [**58**].

In summary, by quantitatively linking signalling levels to motility and interfacial tension, and by showing that invasion is maximized at intermediate adhesion across two mechanical formalisms, we provide a concise physical framework for collective migration in heterogeneous tissues. This framework yields testable predictions for developmental, regenerative, and pathological settings.

## 4 Methods

### 4.1 Simulation setup and non-dimensionalization

Simulations are performed in two dimensions using the vertex energy in Eq. 1. Each cell is represented as a polygon with area *A*_*i*_, perimeter *P*_*i*_, and line tensions *γ*_*ij*_ assigned to each cell–cell edge. Cluster and background tissue cells share the same energy functional and differ through their preferred shape indices, motility parameters, and interfacial line tensions.

All physical units are non-dimensionalized. Lengths are measured in units of the preferred cell size 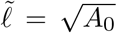 and times in units of 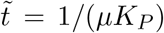, where *µ* is the mobility (inverse friction) in the overdamped dynamics. Throughout the simulations, we set *A*_0_ = 1, *K*_*P*_ = 1, and *µ* = 1, so that the natural units satisfy 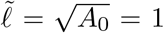 and 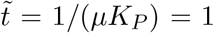. Unless stated otherwise, we use *K*_*A*_ = 10, which keeps cell areas tightly distributed around *A*_0_ even in the presence of large interfacial tensions. Cluster and background mechanics are controlled through their preferred shape indices (equivalently, preferred perimeters), as defined in the main text.

The tissue is simulated in a rectangular domain of size *L*_*x*_×*L*_*y*_ in the (*x, y*) plane, with *L*_*x*_ = *L*_*y*_ = 20 in units of 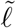. Periodic boundary conditions are imposed along the *y* direction. To mimic an impenetrable edge, an effective rigid wall is imposed on the negative-*x* boundary by pinning the degrees of freedom of cells within a thin strip, preventing migrating cells from crossing this boundary. For comparison with zebrafish transplants [**29**], this boundary can be interpreted as the yolk syncytial layer (YSL), and invasion depth is measured along the *x* axis toward negative *x*.

Initial tissues are generated as disordered confluent configurations from random Voronoi tessellations of uniformly distributed seed points. The system is then relaxed with weak isotropic motility (small *v*_0_ applied uniformly with no directional bias) so that initial Voronoi artefacts are suppressed and the tissue settles into a stable disordered packing with a steady shape-index distribution corresponding to the imposed preferred shape index. Equilibration is run for 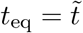, by which time the total energy and cell shapes have converged.

After relaxation, a compact connected group of *N*_*c*_ cells (typically *N*_*c*_ ∈ [1, 8]) is labelled near the positive-*x* side of the domain as the “migrating cluster”, while the remaining cells form the background tissue. Directed motility is then switched on for the active cells of the migrating cluster, and this moment defines *t* = 0. Active forces are biased along the negative-*x* direction so that the cluster migrates toward the rigid boundary. Trajectories are evolved for a measurement window of duration *T* = 100 (in units of 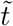), and observables are computed over this interval.

Dynamics are integrated using a forward Euler scheme with time step Δ*t* = 2 × 10^−4^ (in units of 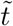). For each parameter combination, *N*_rep_ = 100 independent realisations are simulated with different initial disorder and noise seeds, and summary statistics are computed across these realisations.

All simulations are performed using `CellGPU` [**36**] with custom routines for heterotypic line tensions, active force output, and cluster tracking. Analysis is carried out in Python using standard numerical and graph libraries to process trajectories, assemble connected components, and compute the metrics used above. Code and processed simulation data will be made available upon publication.

### 4.2 Quantification of cluster displacement

Invasion is quantified by the mean cluster displacement along the imposed migration direction. For each cell *i* that belongs to the cluster at *t* = 0, we define the signed displacement

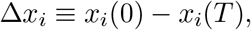

where *T* denotes the end of the measurement window. The mean cluster displacement is then

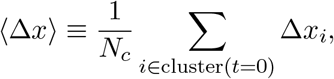

with the sum taken over all cells initially assigned to the cluster.

For mechanically homogeneous tissues, we set 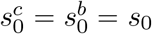 and scan the (*v*_0_, *s*_0_) parameter plane. For mechanically heterogeneous tissues, we vary cluster and background mechanics independently by fixing one shape index and sweeping the other (i.e. fixing 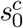 and varying 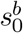, or vice versa) at specified values of *v*_0_. For each parameter combination, ⟨Δ*x*⟩ is averaged over independent realisations, and variability is reported across realisations using either the standard deviation (STD) or the standard error of the mean (SEM), as specified in the corresponding figure legends.

### 4.3 Interfacial coupling: cohesion, shape, and remodelling

Heterotypic interfacial coupling is implemented by assigning a line tension *γ* to all mixed (cluster– background) edges, while homotypic line tensions are set to zero unless stated otherwise. Unless noted, bulk mechanics are fixed to 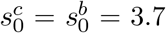 and parameter scans are performed in the (*v*_0_, *γ*) plane for multiple initial cluster sizes *N*_*c*_, using the simulation protocol described above. In addition to the mean displacement ⟨Δ*x*⟩, cluster organisation and interfacial remodelling are quantified as follows.

#### Cluster cohesion (weighted cluster size)

At time *T*, cluster cells are partitioned into connected components based on shared edges. If *n*_*k*_ denotes the number of cells in connected component *k*, the population-weighted cluster size is

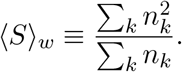

This quantity corresponds to an effective cluster size: ⟨*S*⟩_*w*_ = *N*_*c*_ for a single cohesive cluster and ⟨*S*⟩_*w*_ = 1 for complete dispersion into isolated cells.

#### Cluster shape (circularity)

Cluster shape is quantified for the largest connected component at time *T* . Its outer boundary is used to compute the enclosed area *A*_cl_ and outer perimeter *P*_cl_, defining the circularity

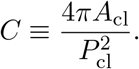

Here *C* = 1 corresponds to a perfect circle and smaller values indicate more elongated or irregular outlines.

#### Interfacial remodelling (front-edge creation and rear-edge detachment)

Remodelling is quantified by the rates at which heterotypic contacts are created at the front and lost at the rear of the cluster. At each saved time point *t*, the cluster centre-of-mass coordinate along the migration axis is computed as *X*_cl_(*t*), and cluster cells are classified as front if *x*_*i*_ > *X*_cl_(*t*) and rear if *x*_*i*_ < *X*_cl_(*t*) (because the migration is in negative *x* direction). Let 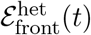 denote the set of heterotypic edges incident to front cluster cells, and 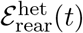 the corresponding set for rear cluster cells. Between successive saved frames separated by Δ*t*_save_, the number of newly formed front heterotypic edges and the number of lost rear heterotypic edges are

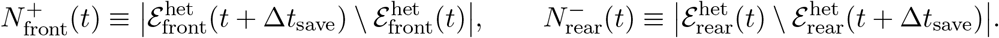

Rates are obtained by normalising by Δ*t*_save_ and by the number of front or rear cells, *N*_front_(*t*) and *N*_rear_(*t*),

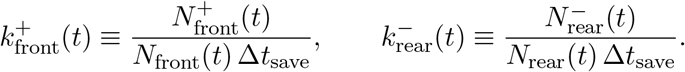

The reported rates are averaged over the measurement window and across independent realisations. Changes in these rates reflect the frequency of local interfacial rearrangements, typically realised via T1 transitions, and are used to identify regimes with active turnover versus kinetically suppressed rearrangements (interfacial pinning).

### 4.4 Leader–follower transport metrics

Leader–follower transport is quantified using two complementary readouts: (i) the displacement of passive (follower) cells and (ii) the leader–follower displacement gap, which quantifies how tightly leaders and followers co-migrate along the migration axis. For each cell *i* of type *α* ∈ {*L, F* } (leaders *L* or followers *F* ) that belongs to the initial cluster at *t* = 0, the signed displacement is

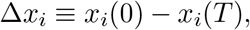

so that Δ*x*_*i*_ > 0 corresponds to net motion along the imposed direction.

Follower transport is measured by the mean follower displacement

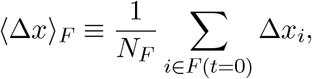

where the sum runs over follower cells in the cluster at *t* = 0 and *N*_*F*_ is the number of such cells.

Leader–follower coordination is quantified by the displacement gap between leaders and followers,

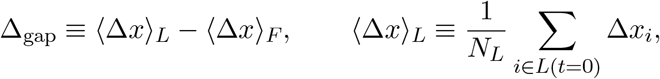

with *N*_*L*_ the number of leader cells at *t* = 0. Small Δ_gap_ indicates coordinated co-migration of leaders and followers, whereas large Δ_gap_ indicates that leaders move ahead of followers, corresponding to weak transport.

### 4.5 Quantification of cluster polarity

Polarity in heterogeneous clusters is quantified by the composition of the front half of the cluster along the migration axis, measured as the fraction of front-half cluster cells that are active (motile). Using the front/rear classification defined above, let 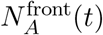 denote the number of active cells with *x*_*i*_ > *X*_cl_(*t*) and let *N* ^front^(*t*) denote the total number of cluster cells in the front half (restricted to cells initially assigned to the cluster at *t* = 0). Cluster polarity is then reported as

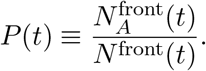

For equal-proportion leader–follower clusters, *P* (*t*) ≃ 0.5 corresponds to a well-mixed configuration, *P* (*t*) = 1 indicates that all active cells are in the front half, and *P* (*t*) = 0 indicates that all active cells are in the rear half (equivalently, follower cells occupy the front half). Time courses are obtained by evaluating *P* (*t*) at each saved time point and averaging across independent realisations.

### 4.6 Fish lines and husbandry

Zebrafish (*Danio rerio*) maintenance and handling were performed using standard procedures [**59, 60**]. The following strains were used in this study: wild-type AB or AB×TL, Tg(*gsc::EGFP-CAAX* ) [**61**], MZoep, and MZoep;Tg(*gsc::EGFP-CAAX* ) [**29**]. Embryos were raised at 25–31^°^C in Danieau’s medium (58 mM NaCl, 0.7 mM KCl, 0.4 mM MgSO_4_, 0.6 mM Ca(NO_3_)_2_, 5 mM HEPES, pH 7.6) and staged according to standard morphological criteria [**60**].

### 4.7 Embryo microinjections

mRNAs were synthesized using the mMessage mMachine kit (Ambion) and injected at the one-cell stage. The following mRNAs were used: 50 pg H2A-chFP [**62**], 50 pg Membrane-RFP [**63**], and 100 pg CA-Mypt [**64**].

### 4.8 Cell transplantation assays

For transplantation assays, donor and host embryos were manually dechorionated with watchmaker forceps in Danieau’s medium. Mesendoderm cells (2–30 cells) were collected from the dorsal margin, marked by EGFP-CAAX expression in Tg(*gsc::EGFP-CAAX* ) embryos, at different stages of gastrulation using a bevelled borosilicate needle (20 *µ*m inner diameter with spike; Biomedical Instruments) connected to a syringe system mounted on a micromanipulator. Donor cells were then transplanted into the dorsal margin of sphere-stage MZoep;Tg(*gsc::EGFP-CAAX* ) host embryos underneath the EVL (schematic representation in Supp. Fig. S7d). The developmental stage of donor embryos is indicated in the corresponding figure panels.

In co-transplantation experiments, mesendoderm cells were sequentially collected from the dorsal margin of donor embryos at different developmental stages into the same bevelled borosilicate needle and then transplanted into the dorsal margin of sphere-stage MZoep;Tg(*gsc::EGFP-CAAX* ) host embryos underneath the EVL (schematic representation in Supp. Fig. S8g). The order of donor-cell collection was purposely varied. This dataset was previously generated in Ref. [**29**] and was reanalyzed here to examine cluster self-organization during internalization.

### 4.9 Image acquisition

For transplantation assays, dechorionated embryos were mounted in a plastic Petri dish containing 2% agarose wells and immobilized in 0.6–0.7% low-melting-point agarose, with the dorsal side of the embryo facing the objective. Embryos were imaged in Danieau’s medium at 28.5^°^C using a Zeiss LSM 880 or LSM 800 upright microscope equipped with a Zeiss Plan-Apochromat 20×/1.0 NA water-immersion objective.

For high-magnification analysis of single-cell shape in MZoep embryos, dechorionated embryos were mounted in a glass-bottom dish (35 mm; MatTek, P35G-1.5-14-C) and immobilized in 0.6–0.7% low-melting-point agarose with the dorsal side of the embryo facing the objective. Embryos were then imaged using a Zeiss LSM 880 inverted microscope equipped with a Zeiss Plan-Apochromat 40×/1.2 NA water-immersion objective.

### 4.10 Experimental Data analysis

Image analysis was performed using Fiji [**65**] and/or Bitplane Imaris.

#### 4.10.1 Analysis of deep cell epiboly

To determine the duration of deep cell epiboly, the Imaris Measurement tool was used to quantify total embryo length and the length covered by the deep cells. Measurements were performed in a single-plane image at the bottom of the imaged volume, as close as possible to the embryo center. The time point at which the deep cells fully covered the yolk cell was defined as 100% epiboly, and the corresponding time was recorded.

#### 4.10.2 Analysis of donor-cell internalization

Donor-cell internalization was quantified as described previously in Ref. [**29**]. Briefly, the Spot plugin of Imaris was used to determine the initial coordinates of donor-cell nuclei, together with reference landmarks at the deep-cell margin. Using a custom MATLAB script [**66**], nuclei and reference landmarks were projected along the *z* axis, and the geometric distance of each projected nucleus to the nearest reference point was computed automatically. Only donor cells positioned within 150 *µ*m of the blastoderm margin of MZoep host embryos were considered for subsequent analysis. Using the same spot-detection workflow aided by cross-sectional projections, both the total number of transplanted donor cells and the number of donor cells that reached the YSL of the host embryo were determined.

### 4.11 Nodal levels used for model parameterization

To compare the model with zebrafish mesendoderm internalization, published measurements of Nodal signalling and transplant behaviour from Ref. [**29**] were used. Effective Nodal levels for donor populations were obtained by reanalyzing the nuclear pSmad2/3 intensity profiles reported at the dorsal margin and restricting the analysis to the first three tiers from the margin (tiers 0–2), corresponding to the mesendoderm progenitor zone from which donor clusters are collected. This yields the discrete signalling levels

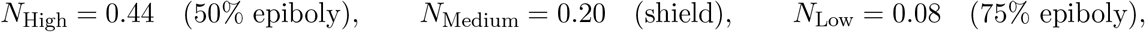

and the host tissue was assigned *N*_host_ = 0 (MZoep background) [**29**]. These values were used as fixed inputs for the Nodal-dependent motility and interaction rules defined above.

### 4.12 Segmentation and cell shape quantification

To parameterize background tissue geometry in vertex-model simulations, membrane-labelled zebrafish ectoderm was imaged by confocal microscopy and projected cell areas and perimeters were extracted from two-dimensional optical sections. For each embryo and time point, a single *z*-plane was selected in which junctional membranes were simultaneously in focus across the analysed patch, yielding a confluent polygonal packing.

Membrane images were intensity-normalized and mildly smoothed by Gaussian filtering prior to segmentation. Cell masks were obtained with Cellpose (v4; `cyto3` model) [**67**] (Supp. Fig. S6a). For each mask, a closed outline was extracted to compute projected area *A* and perimeter *P*, and the per-cell shape index was defined as 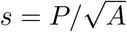.

Geometric filtering was applied to retain only well-resolved bulk ectoderm cell outlines (Supp. Fig. S6b,c). First, a size-based filter removed area outliers to eliminate obvious segmentation artefacts and to exclude very large masks that arise when tissue boundaries enter the field of view (for example, the yolk margin) and are spuriously segmented as single “cells” (Supp. Fig. S6b). Second, a boundary filter excluded non-bulk cells by removing masks at the boundary of the segmented epithelial patch, including partial cells intersecting the image frame, thereby retaining interior cells with complete neighbour contacts for reliable perimeter estimation (Supp. Fig. S6c).

Shape-index distributions were computed from the remaining cells at each time point (Supp. Fig. S6d– g) and summarized by the distribution peak (KDE mode) for each embryo and time point, which is less sensitive than the mean or median to the high-*s* tails introduced by optical sectioning and segmentation artefacts [**68**]. These peak values were then used to compare measured tissue geometry with the corresponding simulated regime.

### 4.13 Zebrafish transplant readouts and best-fit regions

To compare simulations with zebrafish transplant experiments, simulation outcomes are summarised using metrics that mirror the qualitative experimental readouts for homogeneous, mixed, and triple donor compositions.

For a given transplant composition and parameter pair (*α, β*), internalization of a donor subpopulation *C* (e.g. followers) is quantified as the fraction of cells whose displacement along the migration axis exceeds a threshold depth *d*_YSL_,

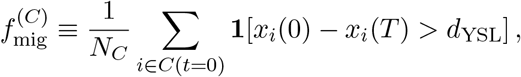

where *C*(*t* = 0) denotes cells of group *C* in the initial donor cluster and **1**[·] is the indicator function. The threshold *d*_YSL_ is chosen from the simulated displacement distributions to match the qualitative notion of successful internalization used in Ref. [**29**].

To assess whether followers remain clustered with leaders, donor cells at time *T* are partitioned into connected components based on shared edges. For a follower group *F* and leader group *L*, the co-clustering fraction is

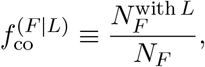

where 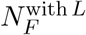 is the number of follower cells that belong to a connected component containing at least one leader cell at time *T* (with *N*_*F*_ counted at *t* = 0). Cluster fragmentation is quantified by a splitting indicator per realisation, equal to 1 if donor cells form two or more connected components at time *T*, and 0 otherwise; the splitting fraction is the proportion of realisations with splitting.

Phase diagrams are constructed by evaluating these metrics on a grid of (*α, β*) for homogeneous High/Medium/Low clusters, mixed High–Medium and High–Low clusters, and triple High–Medium– Low clusters. For each experimental condition, a *best-fit region* in parameter space is defined by threshold criteria chosen to reproduce the reported experimental outcomes in Ref. [**29**] (e.g. strong internalization with limited splitting for homogeneous High; minimal internalization for homogeneous Medium/Low; high co-clustering for High–Medium; low co-clustering and frequent splitting for High– Low; and robust Medium-mediated connectivity in triple transplants without excessive breakup). Best-fit regions are determined using the median of each metric across realizations, and overlap regions identify (*α, β*) pairs that are simultaneously consistent with all experimental constraints for a given adhesion rule.

## Supporting information

SI Theory Note

---

**Fig. S1.**
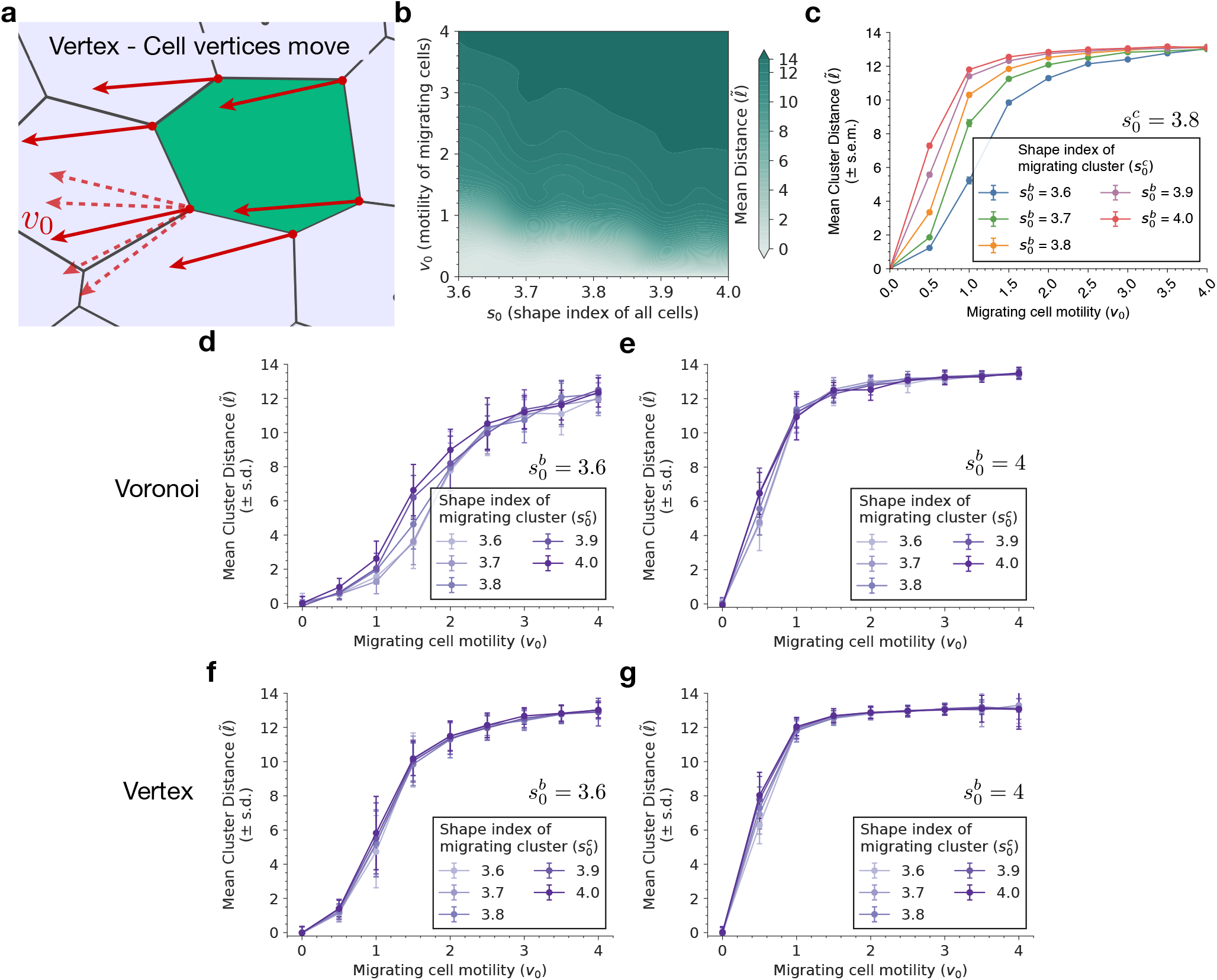
Active Vertex Model (AVM) and comparison with Voronoi simulations. (a-c) AVM setup and migration behavior in homogeneous tissues. Panel (a) shows the vertex-based schematic, in which cell vertices rather than cell centers are evolved under active and mechanical forces. In homogeneous tissues, the mean cluster migration distance increases with motility *v*_0_ and with increasing shape index *s*_0_, as shown by the phase map in (b). At fixed cluster shape index 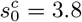, increasing the background shape index 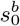 shifts migration to lower motility values, as shown in (c). (d-g) Comparison of the contribution of cluster and background mechanics to invasion, in either SPV or AVM. Panels (d,e) show SPV a results at fixed background shape index 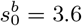 and 4.0, respectively, while varying the cluster shape index 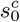. Panels (f,g) show the corresponding AVM parameters. In both models, migration depends primarily on background mechanics, with weaker sensitivity to the cluster shape index.

**Fig. S2.**
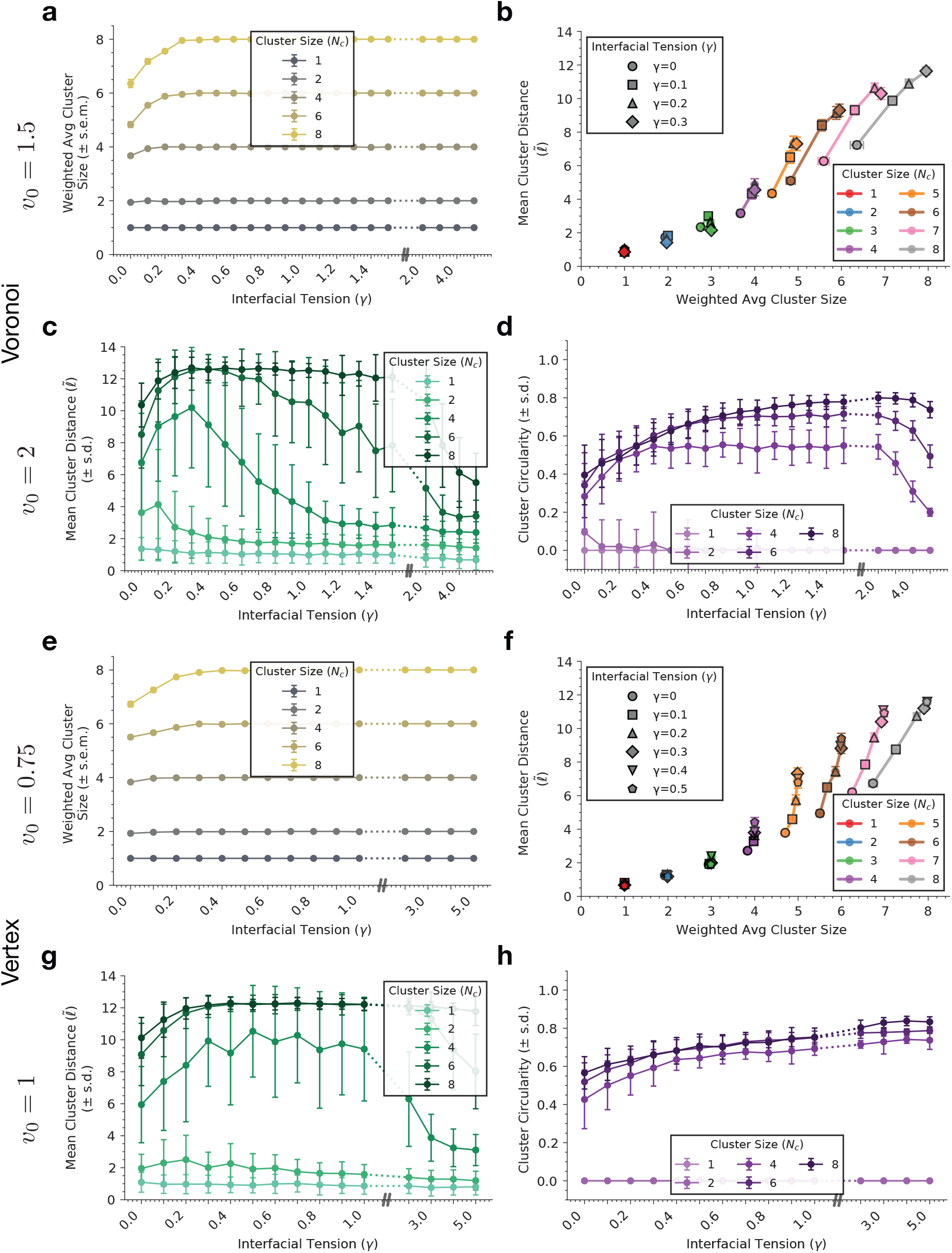
Cluster cohesion, invasion distance, and morphology as a function of heterotypic tension. (a-d) SPV/Voronoi results. Increasing heterotypic tension *γ* rapidly increases the weighted average cluster size until it reaches a plateau, indicating suppression of cluster dispersion (a). Mean cluster distance increases with effective cluster size across the same simulations (b). At higher motility, invasion is maximal at intermediate *γ* and is suppressed at both low and high tension (c). Circularity rises as fragmentation is suppressed and decreases again at large *γ* (d), as discussed in the main text and specifically for the SPV model. (e-h) AVM/Vertex results. As in SPV, increasing *γ* rapidly increases the weighted average cluster size and promotes cohesive clusters (e), and larger effective cluster size is associated with larger invasion distance (f). Mean cluster distance is again maximal at intermediate *γ*, but over a broader range than in SPV (g). In contrast to SPV, circularity remains high at large *γ*, consistent with the distinct high-*γ* morphologies in the AVM (h).

**Fig. S3.**
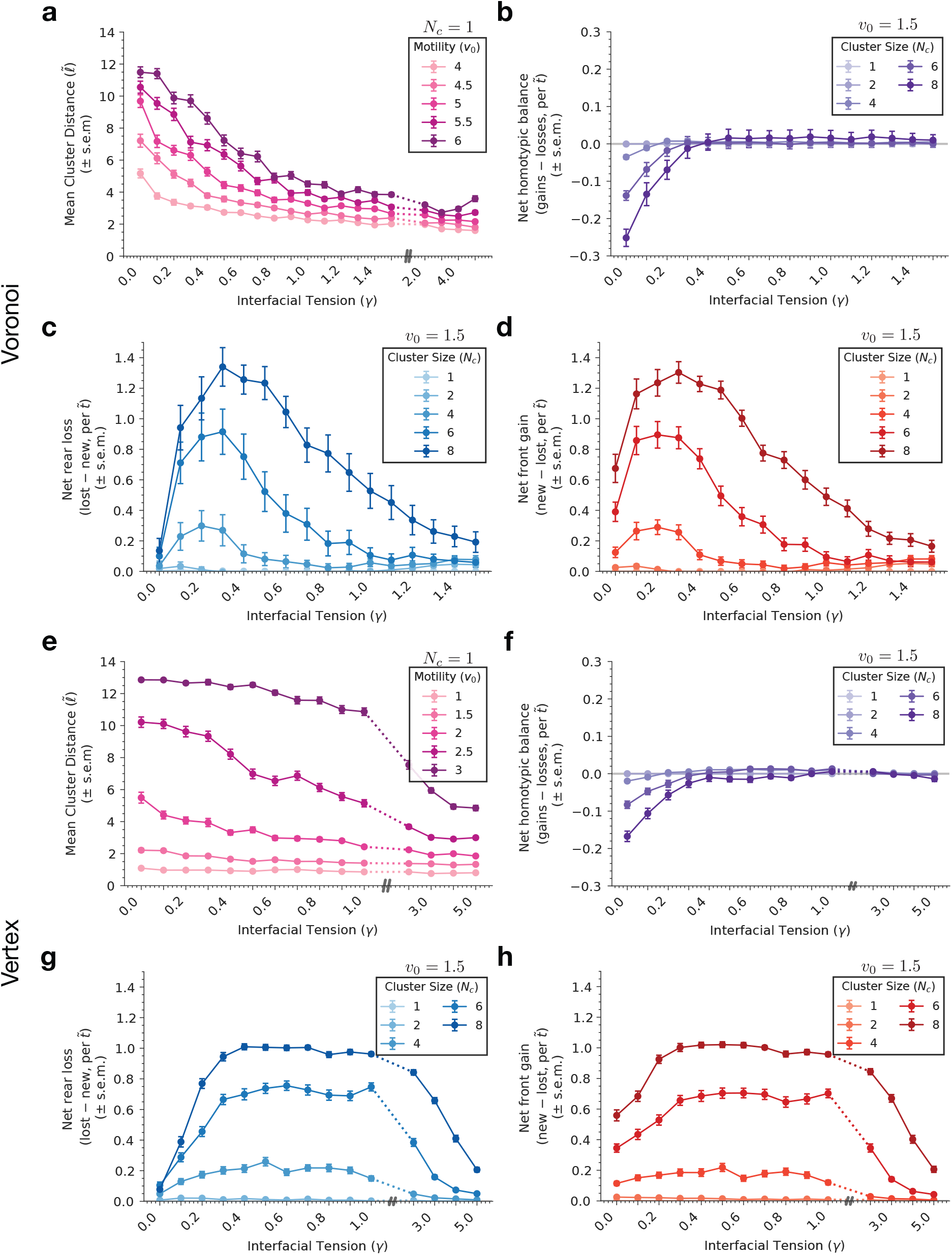
Single-cell migration and interface remodelling as a function heterotypic tension. (a-d) SPV/Voronoi results. For single-cell clusters, invasion decreases monotonously with increasing heterotypic tension *γ* across motility values, indicating that the beneficial effect of *γ* is collective rather than single-cell in origin (a). The net homotypic balance per time step is negative at low *γ* and approaches zero as *γ* increases, reflecting stabilization of within-cluster contacts (b). Net rear-edge loss and net front-edge gain are both maximal at intermediate *γ* and reduced at both low and high tension, consistent with optimal interfacial remodeling in the same regime (c,d). (e-h) AVM/Vertex results. As in SPV, single-cell invasion decreases with increasing *γ* (e), and the net homotypic balance approaches zero as interfacial tension increases (f). Rear-edge loss and front-edge gain are again largest at intermediate *γ*, but over a broader range than in SPV (g,h).

**Fig. S4.**
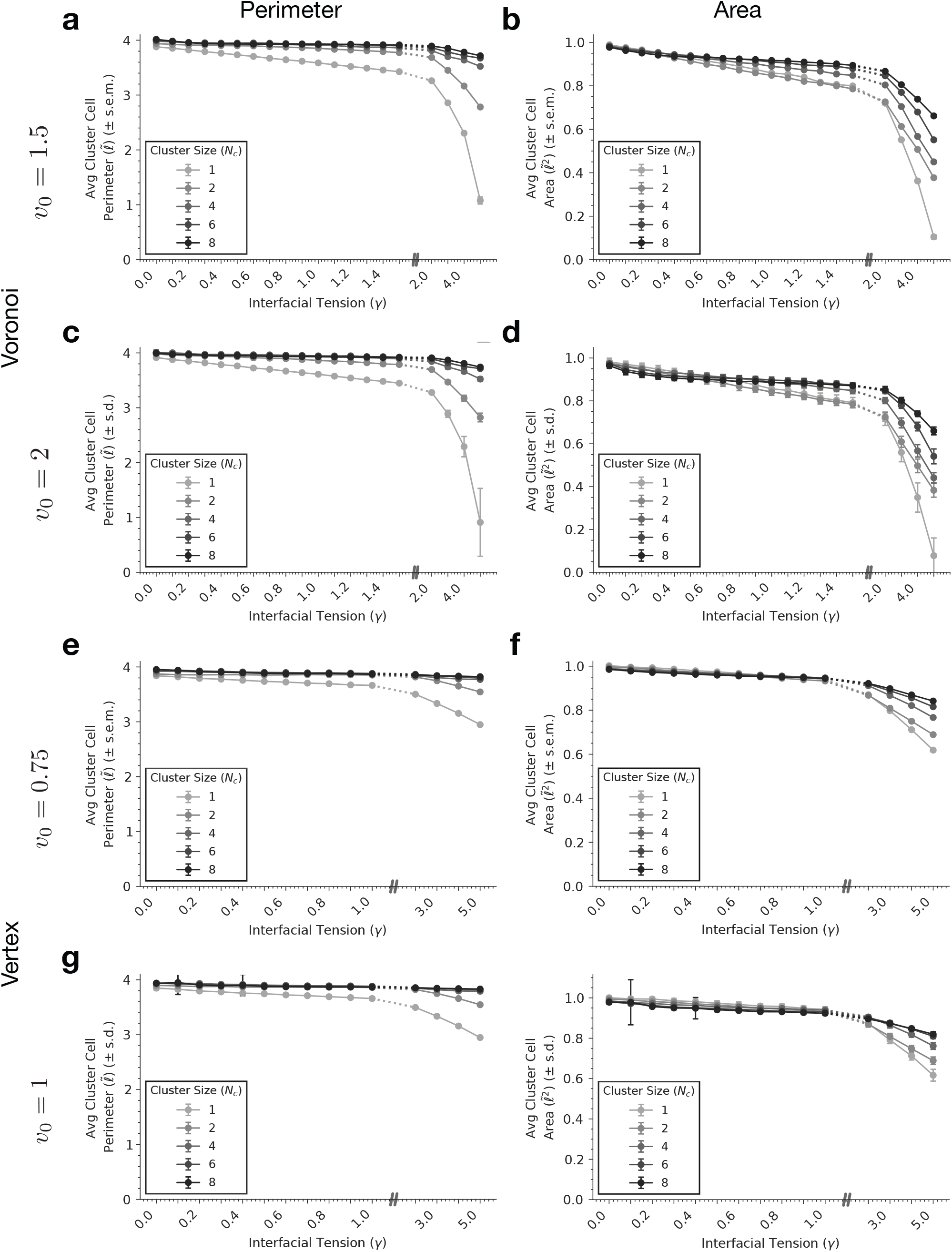
Cluster perimeter and area as a function of heterotypic tension. (a-d) SPV/Voronoi results. Mean cluster-cell perimeter and area are shown as functions of heterotypic tension *γ* at *v*_0_ = 1.5 and *v*_0_ = 2 for different initial cluster sizes *N*_*c*_. In SPV, increasing *γ* reduces both perimeter and area, with a sharper compaction at large *γ* (a-d). (e-h) AVM/Vertex results. Mean cluster-cell perimeter and area are shown as functions of *γ* at *v*_0_ = 0.75 and *v*_0_ = 1 for different initial cluster sizes *N*_*c*_. As in SPV, increasing *γ* compacts the cluster cells and reduces both perimeter and area, although the changes remain more gradual over the range shown (e-h).

**Fig. S5.**
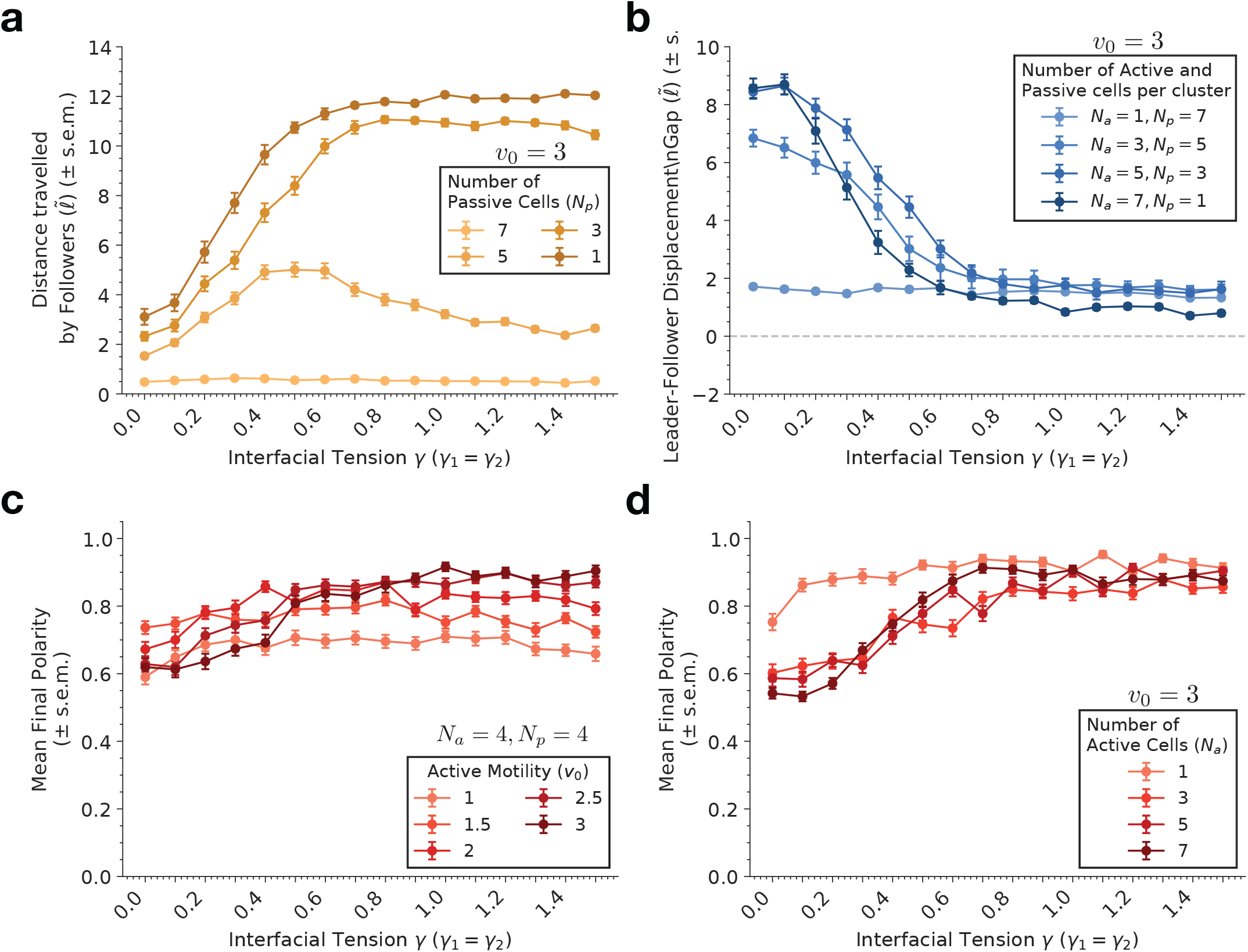
Quantification of leader–follower transport under symmetric heterotypic tension. (a,b) Dependence of leader–follower transport on cluster composition. At fixed active motility *v*_0_ = 3, increasing the number of active cells enhances the distance travelled by passive followers (a) and reduces the leader–follower displacement gap (b). (c,d) Robustness of polarity formation across motility and composition. For fixed leader–follower composition *N*_*a*_ = 4, *N*_*p*_ = 4, the mean final polarity remains high across a broad range of *γ* and *v*_0_ (c). At fixed active motility *v*_0_ = 3, polarity likewise remains robust across different numbers of active cells *N*_*a*_ (d). For this figure, leader–background and follower–background interfacial tensions are set equal, *γ*_1_ = *γ*_2_ *≡ γ*.

**Fig. S6.**
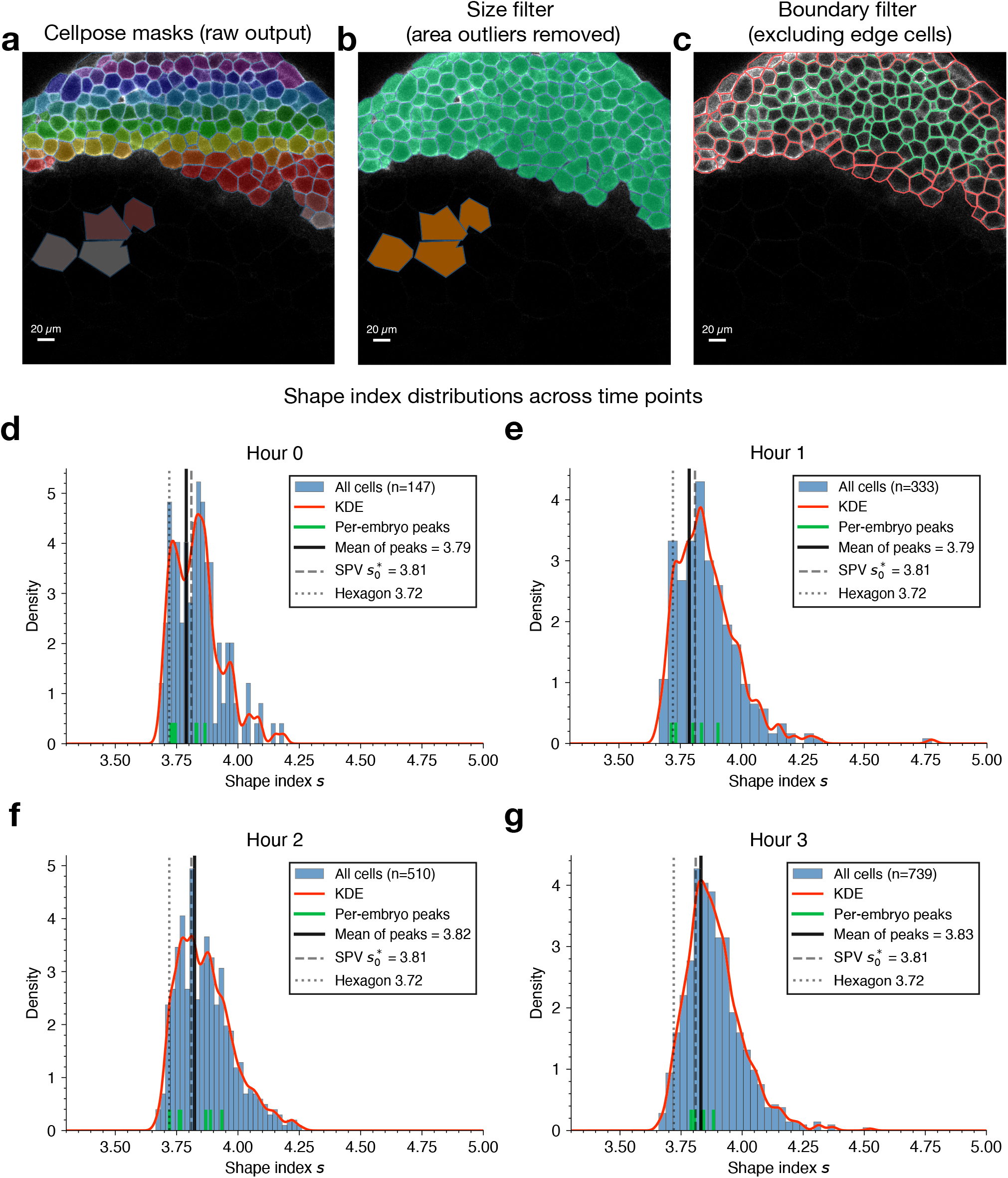
Segmentation and filtering pipeline for estimating the tissue shape index from membrane images. (a-c) Extraction of filtered cell masks from membrane signal images. Panel (a) shows the raw Cellpose output, including incorrectly segmented large cells and other artifacts that do not belong to the blastoderm. Panel (b) shows the mask set after removing area outliers, such as large EVL cells and segmentation artifacts. Panel (c) shows the final boundary-filtered mask (green), excluding edge cells (red), used for shape-index analysis. (d-g) Shape-index distributions across time points. Histograms show the distribution of cell shape index *s* for all filtered cells at hours 0–3 (i.e. the time period at which we assess the migration ability of transplanted cluster cells), together with the corresponding kernel density estimate (red). Green ticks indicate the peak shape index measured independently for each embryo, and the black line marks the mean of these peak values. Across time points, the peak shape index remains close to the SPV reference value 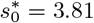 (gray dashed line), and above the regular-hexagon value *s* = 3.72 (gray dotted line).

**Fig. S7.**
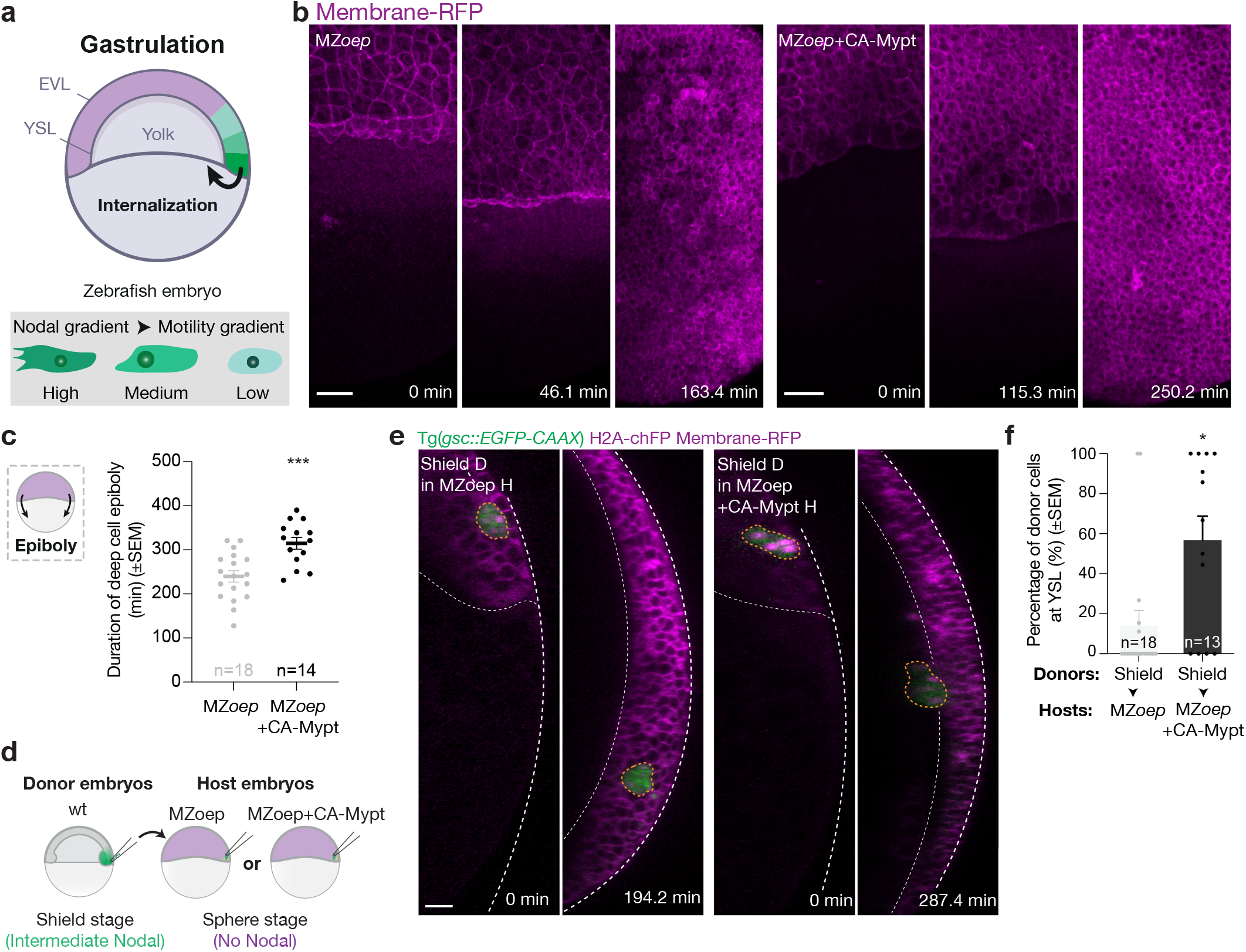
Host-tissue dynamics and internalization of shield-stage donor cells in control and fluidized embryos. (a) Schematic illustrates the Nodal signalling gradient during zebrafish gastrulation, previously shown to encode a gradient of motility forces within the mesendoderm [**29**]. (b) High-resolution confocal images of control and CA-Mypt-injected MZoep embryos expressing Membrane-RFP (magenta) are shown at representative times. (c) Duration of deep cell epiboly is quantified for control and CA-Mypt-injected MZoep host embryos, validating the effect of the CA-Mypt treatment. Data are shown as mean *±* s.e.m.; unpaired *t* test, \*\*\**P* = 0.0004. (d-f) Internalization of shield-stage donor mesendoderm cells in control and fluidized (CA-Mypt) hosts. In (d), a schematic of the transplantation assay is shown. In (e), mesendoderm donor cells collected from the dorsal blastoderm margin of shield-stage embryos were transplanted into control or CA-Mypt-injected MZoep host embryos. Donor cells express Tg(*gsc::EGFP-CAAX* ) (green) and H2A-chFP (magenta, nuclei), while host embryos express low levels of Tg(*gsc::EGFP-CAAX* ) (green) and Membrane-RFP (magenta). For each condition, the first time point (0 min) and the time point at which hosts reached 100% epiboly are shown. In (f), the percentage of shield-stage donor mesendoderm cells located at the YSL at 3 hour time point is quantified for control and CA-Mypt-injected MZoep host embryos. Data are shown as mean *±* s.e.m.; Mann–Whitney test, \**P* = 0.0113. In (e), dashed white lines indicate the EVL and YSL, and dashed outlines mark donor transplants. The dorsal side of the embryo is shown as a top view in (a) and as a cross-section in (d). Scale bars: 50 *µ*m in (b) and 30 *µ*m in (e).

**Fig. S8.**
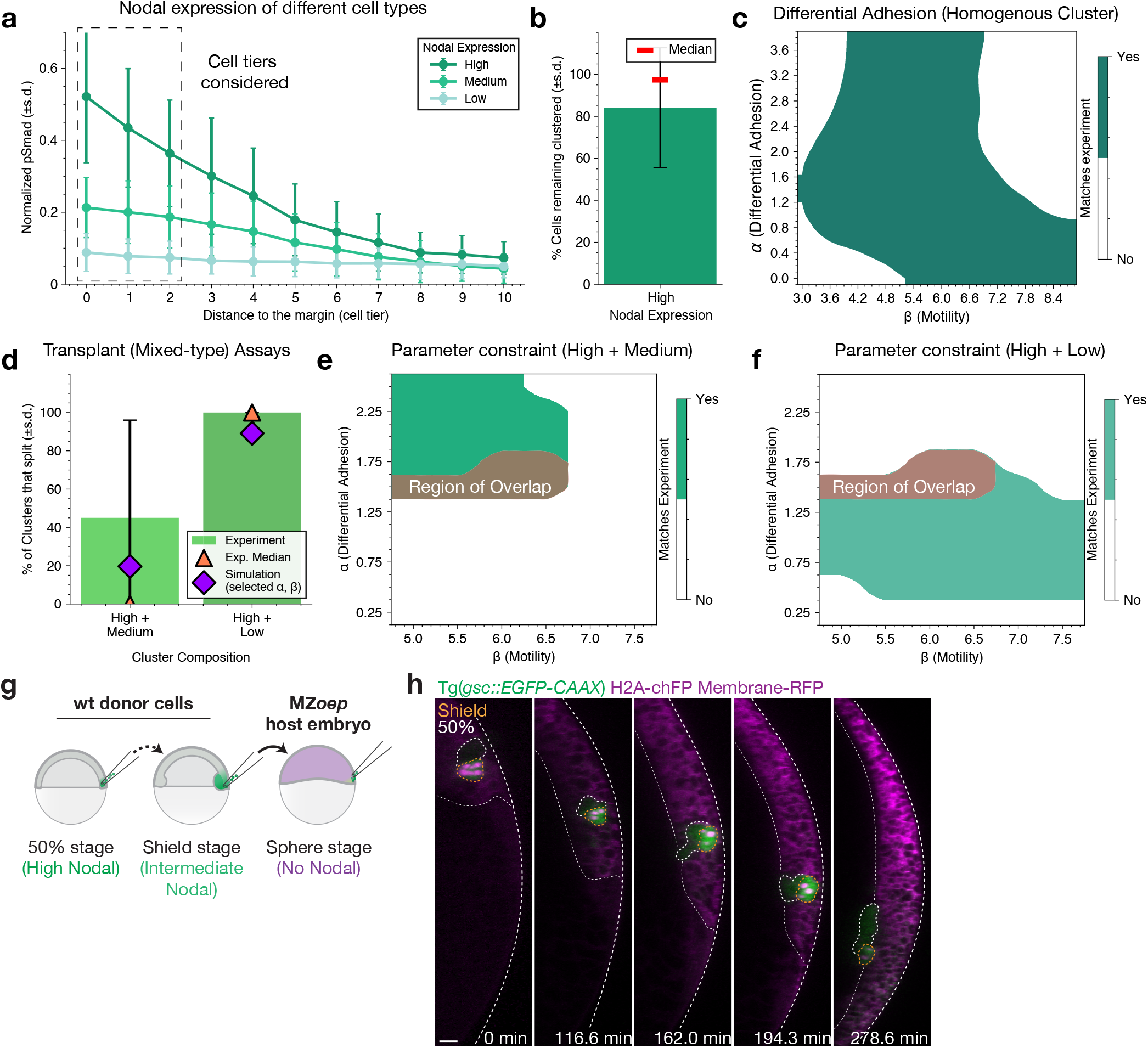
Differential adhesion rule and co-transplant self-organization. (a-c) Parameterization of the differential adhesion rule. In (a), normalized pSmad2/3 levels are shown as a function of distance to the margin for High, Medium, and Low Nodal populations, with the first three cell tiers used to define the effective Nodal levels of clusters, as this is where populations are taken upon transplantation. In (b), the experimental data for the fraction of cells remaining in a single cluster (for High-Nodal cells, i.e. from a 50%-epiboly stage embryo) is shown. In (c), the corresponding parameter constraint in the (*α, β*) plane is shown under the differential adhesion rule, 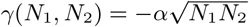. (d-f) Mixed-type transplant assays and model-fitting under the differential adhesion rule. In experiments (d), High+Medium clusters split less frequently than High+Low clusters. Panels (e,f) show the corresponding parameter constraints for the High+Medium and High+Low assays, respectively. The brown region marks the overlap between these two mixed-type constraints. (g,h) Co-transplant assays reveal dynamic leader-follower polarity sorting during collective migration. In (g), a schematic of co-transplanted donor cells collected from 50% epiboly (Nodal-High) and shield-stage (Nodal-Mediaum) embryos is shown. In (h), representative high-resolution confocal images of these co-transplants show that donor cells from 50% epiboly embryos reorganize toward the front of the cluster, whereas shield-stage donor cells reorganize at the rear, illustrating dynamical sorting during invasion. In (h), dashed white lines indicate the EVL and YSL, and dashed outlines mark donor transplants. The dorsal side of the embryo is shown as a cross-section in (g). Scale bar: 30 *µ*m in (h).

**Fig. S9.**
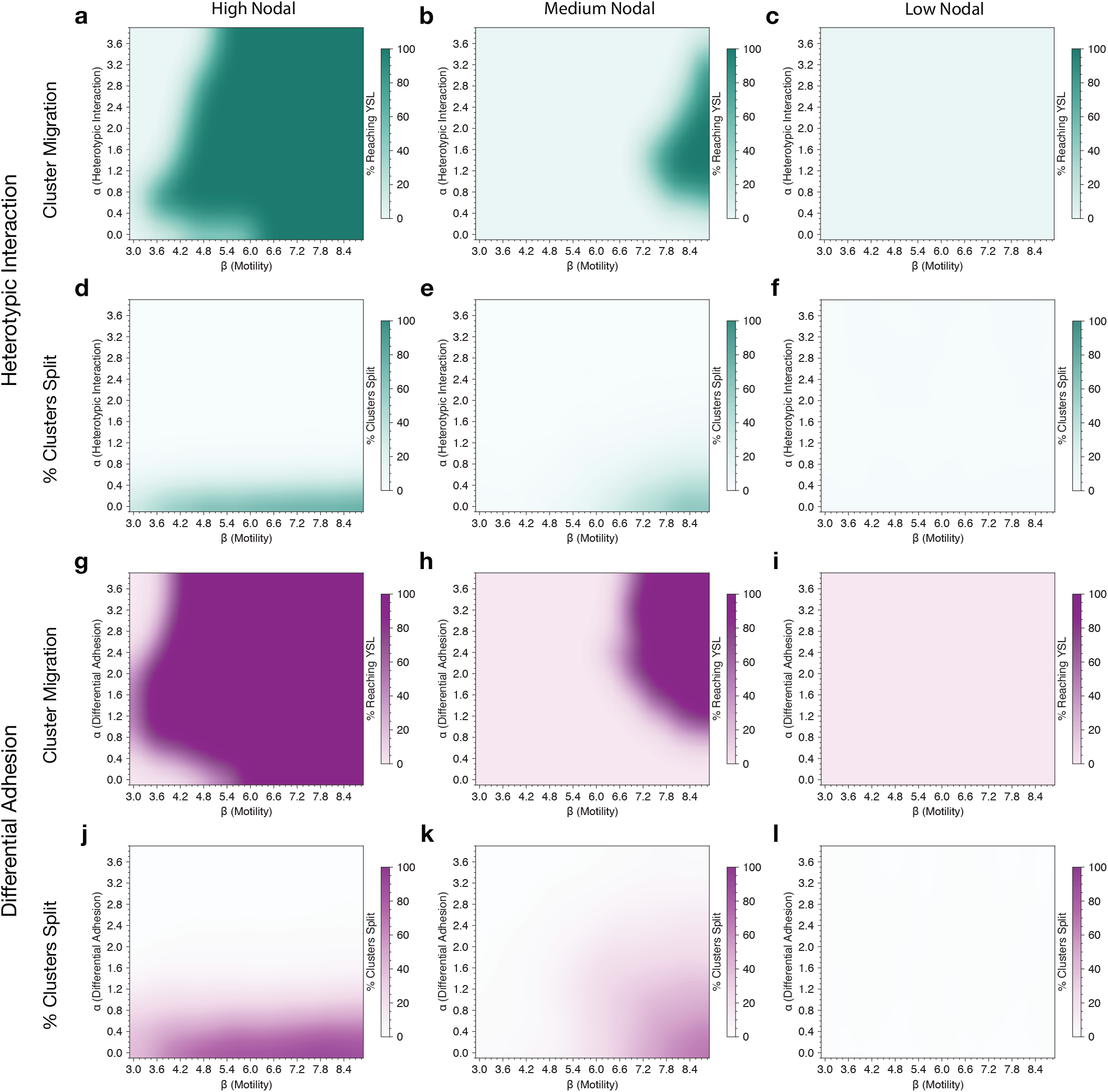
Homogeneous cluster migration and splitting under heterotypic and differential adhesion rules. (a-f) Phase diagram of simulation outputs under heterotypic adhesion rule. Panels (a-c) show the fraction of donor cells that reach the YSL threshold for High, Medium, and Low Nodal clusters, respectively. Panels (d-f) show the corresponding fraction of clusters that split into disconnected pieces by the final time point. Together, these maps identify parameter regimes in which High-Nodal clusters migrate efficiently whereas Medium and Low-Nodal clusters remain largely non-migratory and intact. (g-l) Phase diagram of simulation outputs under the differential adhesion rule. Panels (g-i) show the fraction of donor cells that reach the YSL threshold for High, Medium, and Low Nodal clusters, respectively. Panels (j-l) show the corresponding fraction of clusters that split into disconnected pieces by the final time point. As in the heterotypic case, these maps define the homogeneous-cluster constraints used in the main-text parameter scans. In all panels, color indicates the median fraction across simulations at each (*α, β*).

**Fig. S10.**
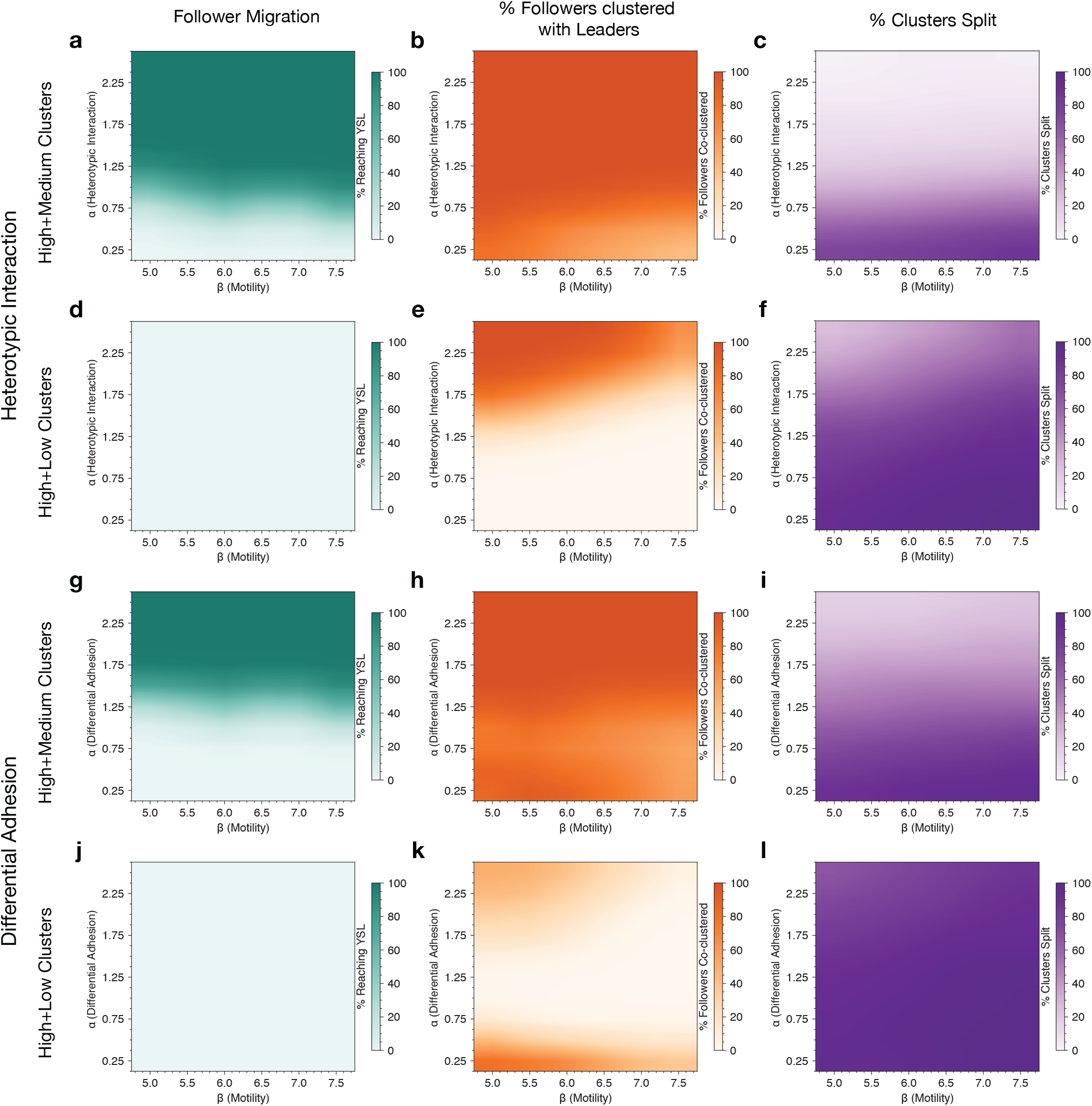
Two-cell-type cluster migration, co-clustering, and splitting under heterotypic interaction and differential adhesion rules. (a-f) Phase diagram of simulation outputs under the heterotypic interaction rule. Panels (a-c) show the follower migration fraction, follower co-clustering fraction with leaders, and cluster-splitting fraction, respectively, for High–Medium clusters. Panels (d-f) show the corresponding quantities for High–Low clusters. Under the heterotypic rule, High–Medium clusters combine efficient follower transport with strong co-clustering, whereas High–Low clusters show weak follower migration, reduced co-clustering, and increased splitting. (g-l) Phase diagram of simulation outputs under the differential adhesion rule. Panels (g-i) show the follower migration fraction, follower co-clustering fraction with leaders, and cluster-splitting fraction, respectively, for High–Medium clusters. Panels (j-l) show the corresponding quantities for High–Low clusters. Under the differential rule, High–Medium clusters can still migrate collectively, but High–Low clusters exhibit weak follower transport, reduced co-clustering, and extensive splitting. In all panels, color indicates the median value across simulations at each (*α, β*). These maps provide the mixed-cluster agreement masks used in Fig. 4 and Fig. S8.

**Fig. S11.**
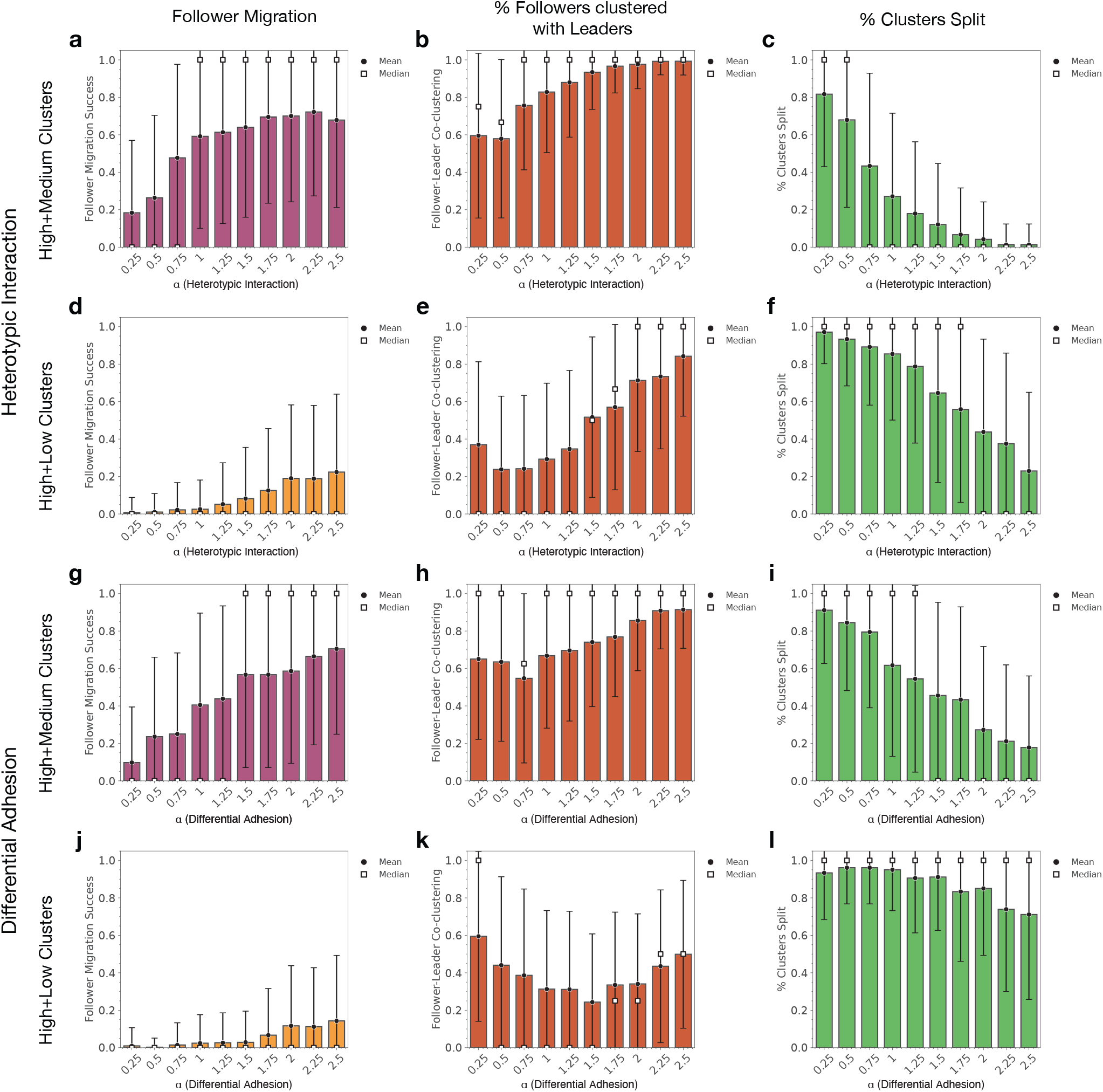
Mixed-cluster metric distributions at fixed motility support the use of median values. All panels show simulations at fixed motility *β* = 5.5 as a function of adhesion strength *α*. Bars indicate the distribution across simulations for each condition, while circles and squares denote the mean and median, respectively. (a-f) Simulation results under heterotypic interaction rule. Panels (a-c) show follower migration success, follower co-clustering with leaders, and cluster-splitting fraction, respectively, for High–Medium clusters. Panels (d-f) show the corresponding quantities for High–Low clusters. Broad and often skewed distributions lead to noticeable differences between mean and median values. (g-l) Simulation results under differential adhesion rule. Panels (g-i) show follower migration success, follower co-clustering with leaders, and cluster-splitting fraction, respectively, for High–Medium clusters. Panels (j-l) show the corresponding quantities for High–Low clusters. As in the heterotypic case, substantial variability across simulations produces systematic differences between mean and median values. These distributions motivate the use of median-based agreement masks in Fig. 4 and Supp. Figs. S8–S10, which are more robust to outliers than mean-based summaries.

**Fig. S12.**
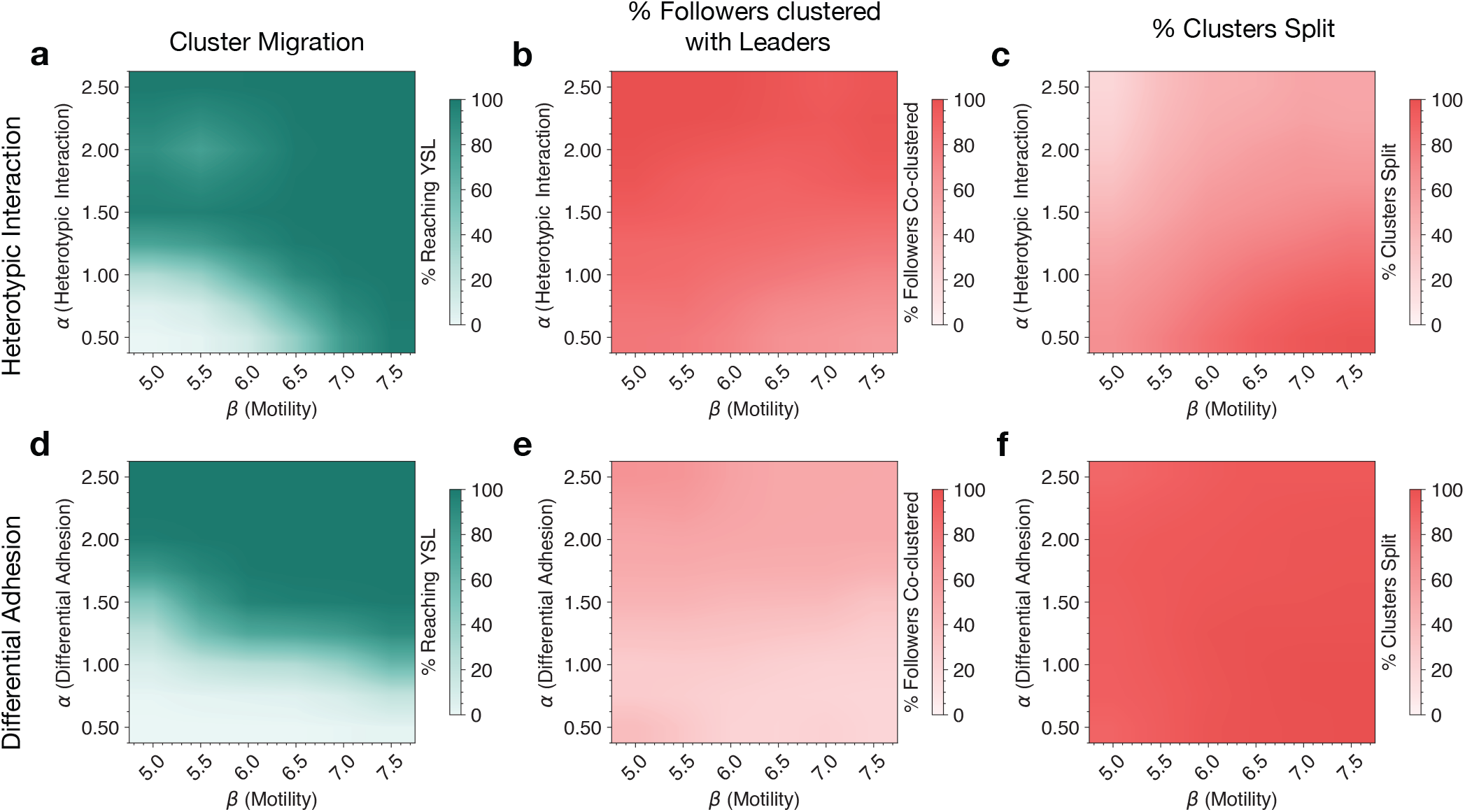
Three-cell-type cluster migration, co-clustering, and splitting under heterotypic interaction and differential adhesion rules. (a-c) Simulation results under the heterotypic interaction rule. Panels (a-c) show the cluster migration fraction, follower co-clustering fraction, and cluster-splitting fraction, respectively, for clusters containing High, Medium, and Low Nodal cells. Under the heterotypic rule, collective migration is accompanied by strong co-clustering and comparatively limited splitting over a substantial region of parameter space. (d-f) Simulation results under the differential adhesion rule. Panels (d-f) show the same three summary metrics for the same three-cell-type clusters. Although substantial cluster migration can still occur, it is associated with weaker co-clustering and extensive splitting, indicating selective dispersion rather than cohesive collective migration. In all panels, color indicates the median value across simulations at each (*α, β*). These maps define the parameter ranges used to interpret the three-cell-type assay in Fig. 5.

## Notes

### Competing Interest Statement

The authors have declared no competing interest.

### Funding

- European Research Council: 101116586

