## Supplementary material for "Logic of optimal collective migration in heterogeneous tissues": SI Theory Note

### 1 Supplementary Note 1: Vertex-model formulations and interfacial line tension

#### 1.1 Vertex-model formulations

We model migrating clusters in confluent tissues using the same two-dimensional vertex framework as in the main text, implemented in two complementary realizations: the Self-Propelled Voronoi (SPV) model and the Active Vertex Model (AVM) [1–4]. In both cases, each cell is represented by a polygon with area $A_i$ and perimeter $P_i$, while neighboring cells $i$ and $j$ share an interface of length $l_{ij}$ (see Extended Fig. 1 and Extended Fig. 2 for the two geometric realizations, and Fig. 1 in the main text for the biological setup).

The mechanical energy is

$$E = \sum_i \Big[ K_A (A_i - A_0)^2 + K_P (P_i - P_0)^2 \Big] + \sum_{\langle i,j \rangle} \gamma_{ij}\, l_{ij}, \tag{1}$$

where $K_A$ and $K_P$ are the stiffnesses associated with area and perimeter deviations, and $\gamma_{ij}$ denotes the line tension assigned to the edge shared by cells $i$ and $j$. As in the main text, each cell is labeled by its type $\sigma_i \in \{c, b\}$, corresponding to cluster and background cells, respectively, and we define

$$\gamma_{ij} = \begin{cases} \gamma_{\text{het}}, & \text{if } \sigma_i \neq \sigma_j \quad \text{(heterotypic edge, cluster–host interface)}, \\ \gamma^{(c)}_{\text{hom}}, & \text{if } \sigma_i = \sigma_j = c \quad \text{(cluster–cluster edge)}, \\ 0, & \text{if } \sigma_i = \sigma_j = b \quad \text{(background–background edge)}. \end{cases} \tag{2}$$

Unless otherwise stated, we set $\gamma^{(c)}_{\text{hom}} = 0$ and take background–background tensions to vanish, varying only $\gamma_{\text{het}}$.

The dynamics are overdamped. Denoting by $\mathbf{x}_i$ the relevant degrees of freedom, the equations of motion take the generic form

$$\dot{\mathbf{x}}_i = -\,\mu\, \nabla_{\mathbf{x}_i} E + \mathbf{v}_i, \tag{3}$$

where $\mu$ is the mobility and $\mathbf{v}_i$ is the active migration term. Following the main text, we model $\mathbf{v}_i$ as a persistent active motility term of typical magnitude $v_0$, written as

$$\mathbf{v}_i = v_0 \mathbf{n}_i, \qquad \mathbf{n}_i = (\cos\theta_i, \sin\theta_i), \tag{4}$$

with polarity angle $\theta_i$ evolving as an Ornstein–Uhlenbeck process with persistence time $\tau$,

$$\dot{\theta}_i = \frac{1}{\tau}\big(\phi_i - \theta_i\big) + \sqrt{2D_r}\, \eta_i(t), \tag{5}$$

where $D_r$ is the rotational diffusion coefficient and $\eta_i(t)$ is a Gaussian white noise with zero mean and unit variance. In the invasion simulations, the preferred direction $\phi_i$ introduces a weak bias along the $-x$ direction, consistent with the guided migration described in the main text. Polarity vectors are taken to fluctuate independently across neighboring cells.

As in the main text, the equations are non-dimensionalized by setting $A_0 = 1$, which fixes the length scale, and by measuring time in units of $1/(\mu K_P)$, so that $\mu$ can be set to 1 without loss of generality. The preferred perimeter is then equivalently parameterized by the target shape index $s_0 = P_0/\sqrt{A_0}$. Unless stated otherwise, we use $K_A = 10$ and $K_P = 1$, which keeps cell areas close to their targets while allowing shape fluctuations and neighbor exchanges. After this rescaling, we keep the same notation as in the main text for all variables and parameters.

The same mechanical energy is realized differently in the two model variants, depending on the chosen degrees of freedom. In SPV, the dynamical variables are the cell centers $\{\mathbf{r}_i\}$, from which the Voronoi tessellation is constructed (Extended Fig. 1). Cell areas, perimeters, and interface lengths are obtained directly from this tessellation, and topological rearrangements arise naturally through updates of the dual Delaunay triangulation. In AVM, the dynamical variables are the polygon vertices $\{\boldsymbol{\nu}\}$, and cells are represented explicitly as polygons (Extended Fig. 2). In this case, forces act directly on vertices and topological transitions are implemented when edges become sufficiently short. The following subsections describe these two realizations in detail.

##### 1.1.1 Self-Propelled Voronoi (SPV)

In the Voronoi realization, the degrees of freedom are the cell centers $\{\mathbf{r}_i\}$ that generate the tessellation shown in Extended Fig. 1. A small displacement of a center modifies the local Delaunay triangulation and its dual Voronoi cells, so areas, perimeters, and interface lengths are obtained directly from the instantaneous geometry. Neighbor exchanges (T1 transitions) arise naturally: when four cell centers become locally co-circular, the shared Delaunay edge is flipped and the corresponding Voronoi edge is reassigned, updating the neighbor graph without requiring an additional rule.

Cell centers evolve according to the overdamped active dynamics introduced above [Eq. (3)], where $\mathbf{r}_i$ denotes the center position and $\mathbf{n}_i = (\cos\theta_i, \sin\theta_i)$ the instantaneous polarity direction.

The mechanical force on cell $i$ is written as

$$\mathbf{F}_i = -\nabla_{\mathbf{r}_i} E = \mathbf{F}_i^{\text{bulk}} + \mathbf{F}_i^{\text{tens}},$$

separating bulk and interfacial contributions. Both parts are organized through the Voronoi vertices $\mathbf{h}_{ijk}$ associated with Delaunay triangles $(i, j, k)$,

$$\mathbf{F}_i = -\sum_{(i,j,k)} \left[\frac{\partial E_{\text{bulk}}}{\partial \mathbf{h}_{ijk}} + \frac{\partial E_{\text{tens}}}{\partial \mathbf{h}_{ijk}}\right]^{\top} \frac{\partial \mathbf{h}_{ijk}}{\partial \mathbf{r}_i}, \tag{6}$$

which mirrors the geometric construction in Extended Fig. 1. The terms $\partial E_{\text{bulk}}/\partial \mathbf{h}_{ijk}$ and $\partial E_{\text{tens}}/\partial \mathbf{h}_{ijk}$ describe how area, perimeter, and interface energies vary when a single Voronoi vertex moves, while the Jacobian $\partial \mathbf{h}_{ijk}/\partial \mathbf{r}_i$ maps this local vertex response to the motion of the cell center.

Combining these elements yields the compact force expressions used in simulations,

$$\mathbf{F}_i^{\text{bulk}} = -\sum_{(i,j,k)} \left( \frac{\partial \mathbf{h}_{ijk}}{\partial \mathbf{r}_i} \right)^{\top} \sum_{c \in \{i,j,k\}} \left[ 2K_A(A_c - A_0) \frac{\partial A_c}{\partial \mathbf{h}_{ijk}} + 2K_P(P_c - P_0) \frac{\partial P_c}{\partial \mathbf{h}_{ijk}} \right], \tag{7}$$

$$\mathbf{F}_i^{\text{tens}} = \sum_{(i,j,k)} \left( \frac{\partial \mathbf{h}_{ijk}}{\partial \mathbf{r}_i} \right)^{\top} \sum_{(a,b) \in \{(i,j),(j,k),(k,i)\}} \gamma_{ab} \, \hat{\mathbf{t}}^{(ab)}. \tag{8}$$

Here, $\gamma_{ab}$ is the line tension between cells $a$ and $b$, and $\hat{\mathbf{t}}^{(ab)}$ is the unit tangent along their shared edge oriented away from the vertex. All geometric derivatives $\partial A_c/\partial \mathbf{h}_{ijk}$, $\partial P_c/\partial \mathbf{h}_{ijk}$, and $\partial \mathbf{h}_{ijk}/\partial \mathbf{r}_i$ are derived in Appendix A (Eqs. (24)–(31)).

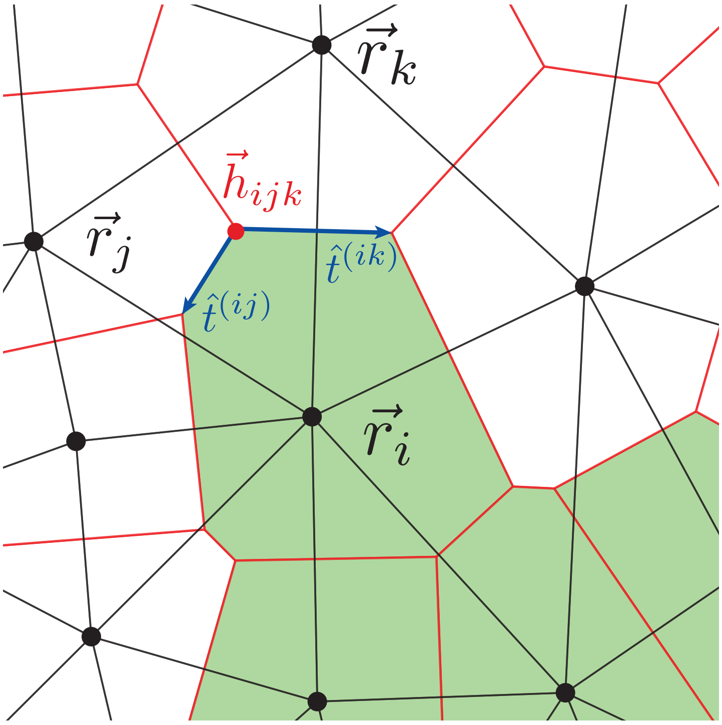

**Extended Fig. 1: SPV geometry.** Cell center positions are specified by vectors $\mathbf{r}_i$ (black dots). They form a Delaunay triangulation (black lines). Its dual is the Voronoi tessellation (red lines), with vertices $\mathbf{h}_{ijk}$ (red dot). Shaded regions indicate different cell types, with cell $i$ highlighted. Unit tangents $\hat{\mathbf{t}}^{(ij)}$ and $\hat{\mathbf{t}}^{(ik)}$ (blue arrows) are oriented away from the Voronoi vertex.

##### 1.1.2 Active Vertex Model (AVM)

In the vertex realization, the degrees of freedom are the polygon vertices $\{\boldsymbol{\nu}\}$ that define the cell boundaries, as shown in Extended Fig. 2. Each vertex is shared by up to three cells, and all geometric quantities such as areas, perimeters, and edge lengths are obtained directly from the polygonal mesh. A small vertex displacement locally changes the geometry of the neighboring cells. Topological transitions (T1) are introduced explicitly: when an edge shortens below a threshold length $\ell_{\min}$, the shared edge is reassigned to connect the opposite pair of cells, updating the neighborhood configuration.

Vertices follow the same overdamped dynamics introduced earlier [Eq. (3)], where $\boldsymbol{\nu}$ denotes the vertex position. Each cell carries a polarity vector $\mathbf{n}_i = (\cos\theta_i, \sin\theta_i)$, and in AVM the active drive at a vertex is taken to be the average of the propulsive contributions of the cells incident on that vertex,

$$\mathbf{v}_{\text{act}}(\boldsymbol{\nu}) = \frac{v_0}{3} \sum_{i\in\mathcal{N}(\boldsymbol{\nu})} \mathbf{n}_i,$$

where $\mathcal{N}(\boldsymbol{\nu})$ denotes the set of cells meeting at $\boldsymbol{\nu}$. This provides a simple vertex-level representation of cell-based propulsion.

The mechanical force acting on a vertex is written as

$$\mathbf{F}(\boldsymbol{\nu}) = -\nabla_{\boldsymbol{\nu}} E = \mathbf{F}_{\text{bulk}}(\boldsymbol{\nu}) + \mathbf{F}_{\text{tens}}(\boldsymbol{\nu}),$$

separating the bulk and interfacial contributions. Both parts are organized through the three cells $(i, j, k)$ that meet at each vertex,

$$\mathbf{F}(\boldsymbol{\nu}) = -\left[\frac{\partial E_{\text{bulk}}}{\partial \boldsymbol{\nu}} + \frac{\partial E_{\text{tens}}}{\partial \boldsymbol{\nu}}\right],$$

which mirrors the geometric construction in Extended Fig. 2. The first term captures how area and perimeter energies change when the vertex moves, while the second term collects the effect of line tension along the adjoining interfaces.

Combining these elements gives the compact force expressions used in simulations,

$$\mathbf{F}_{\text{bulk}}(\boldsymbol{\nu}) = -\sum_{c\in\{i,j,k\}} \left[2K_A(A_c - A_0)\frac{\partial A_c}{\partial \boldsymbol{\nu}} + 2K_P(P_c - P_0)\frac{\partial P_c}{\partial \boldsymbol{\nu}}\right], \tag{9}$$

$$\mathbf{F}_{\text{tens}}(\boldsymbol{\nu}) = \sum_{(a,b)\in\{(i,j),(j,k),(k,i)\}} \gamma_{ab}\,\hat{\mathbf{t}}^{(ab)}. \tag{10}$$

Here, $\gamma_{ab}$ is the line tension between cells $a$ and $b$, and $\hat{\mathbf{t}}^{(ab)}$ is the unit tangent along their shared edge oriented away from the vertex. All geometric derivatives $\partial A_c/\partial\boldsymbol{\nu}$ and $\partial P_c/\partial\boldsymbol{\nu}$ are provided in Appendix B (Eqs. (35)–(40)). These forces are evaluated directly from the current geometry, and the vertex positions are then updated according to Eq. (3) to obtain the AVM dynamics.

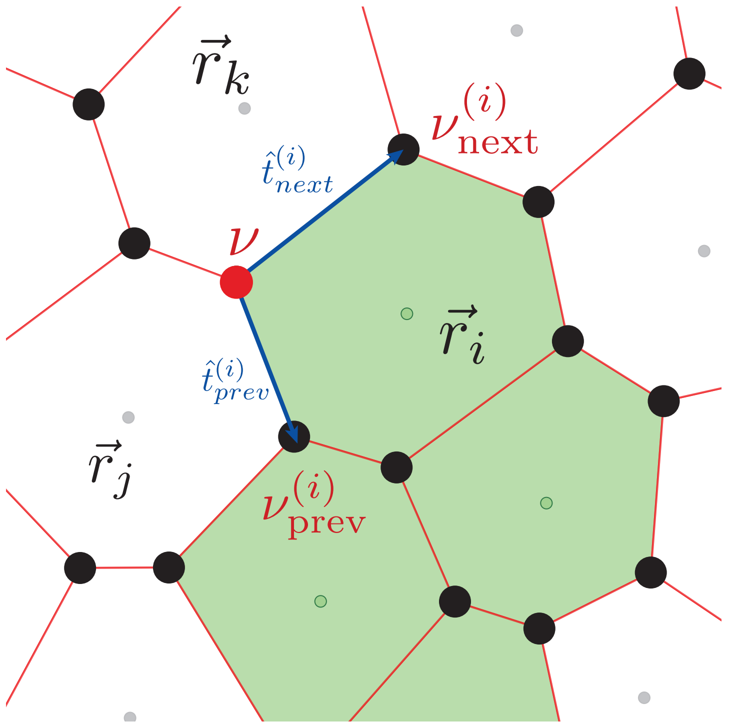

**Extended Fig. 2: AVM geometry.** Cells are polygons with vertices $\boldsymbol{\nu}$ (black dots). Shaded regions indicate different cell types; shown are cells $i$ and $j$ with centers $\mathbf{r}_i$ and $\mathbf{r}_j$. At the focal vertex $\boldsymbol{\nu}$ (red dot), adjacent vertices $\boldsymbol{\nu}^{(i)}_{\text{prev}}$ and $\boldsymbol{\nu}^{(i)}_{\text{next}}$ define the local edges of cell $i$. Unit tangents $\hat{\mathbf{t}}^{(i)}_{\text{prev}}$ and $\hat{\mathbf{t}}^{(i)}_{\text{next}}$ (blue arrows) are oriented away from $\boldsymbol{\nu}$ along these edges.

##### 1.1.3 Geometric basis of the SPV–AVM response differences

The essential distinction between SPV and AVM formulations lies in their choice of degrees of freedom and in how mechanical responses propagate through the tissue. In SPV, the independent coordinates are the cell centers $\{\mathbf{r}_i\}$, which define the Voronoi tessellation through circumcenters $\mathbf{h}_{ijk}$. In AVM, they are the polygon vertices $\{\boldsymbol{\nu}\}$, which directly delineate cell boundaries. This geometric choice determines how local shape changes translate into mechanical forces and thereby leads to distinct migration modes.

*SPV: distributed response.* A displacement of a cell center $\delta\mathbf{r}_i$ perturbs all Delaunay triangles sharing $i$. Since each Voronoi vertex $\mathbf{h}_{ijk}$ depends on three centers,

$$\delta\mathbf{h}_{ijk} = \frac{\partial\mathbf{h}_{ijk}}{\partial\mathbf{r}_i}\,\delta\mathbf{r}_i + \frac{\partial\mathbf{h}_{ijk}}{\partial\mathbf{r}_j}\,\delta\mathbf{r}_j + \frac{\partial\mathbf{h}_{ijk}}{\partial\mathbf{r}_k}\,\delta\mathbf{r}_k, \tag{11}$$

where $\partial\mathbf{h}_{ijk}/\partial\mathbf{r}_i$ is the Jacobian of the circumcenter with respect to its generating centers. Area and perimeter variations follow as

$$\delta A_i = \sum_{(i,j,k)} \frac{\partial A_i}{\partial\mathbf{h}_{ijk}} \cdot \frac{\partial\mathbf{h}_{ijk}}{\partial\mathbf{r}_i}\,\delta\mathbf{r}_i, \qquad \delta P_i = \sum_{(i,j,k)} \frac{\partial P_i}{\partial\mathbf{h}_{ijk}} \cdot \frac{\partial\mathbf{h}_{ijk}}{\partial\mathbf{r}_i}\,\delta\mathbf{r}_i. \tag{12}$$

Each $\delta\mathbf{r}_i$ therefore induces correlated deformations in multiple adjacent cells. The corresponding force variation is

$$\delta\mathbf{F}_i = -\frac{\partial(\nabla_{\mathbf{r}_i} E)}{\partial \mathbf{r}_i}\,\delta\mathbf{r}_i,$$

which includes mixed terms with neighbors and produces a more distributed geometric coupling across the local tissue neighborhood.

*AVM: local response.* In AVM, a displacement of a vertex $\delta\boldsymbol{\nu}$ affects only the three cells meeting at that vertex:

$$\delta A_i = \frac{\partial A_i}{\partial \boldsymbol{\nu}} \cdot \delta\boldsymbol{\nu}, \qquad \delta P_i = \frac{\partial P_i}{\partial \boldsymbol{\nu}} \cdot \delta\boldsymbol{\nu}. \tag{13}$$

The resulting mechanical response therefore remains local: a vertex displacement modifies only the areas, perimeters, and edge lengths of the incident cells and interfaces, in contrast to the more distributed geometric response in SPV.

*Implications for migration.* Both realizations follow Eq. (3), so motility enters as an additive active drive with magnitude $v_0$. The difference lies in how this drive is transmitted through the tissue geometry. In SPV, a displacement of a cell center modifies several neighboring Voronoi vertices through the circumcenter Jacobians, producing a more distributed mechanical response. In AVM, the same active forcing acts through vertex displacements and therefore remains localized to the incident cells and interfaces. These differences in geometric coupling provide a natural basis for the distinct migration modes observed in the two formulations. Algebraic details are provided in Appendices A and B.

#### 1.2 Line-tension conventions and effective perimeter renormalization

In the main text, heterotypic line tension is introduced as the key control parameter for cluster–background interfaces. Here we only clarify a complementary technical point: a spatially uniform shift in line tension can be absorbed into the preferred perimeter, so that the mechanically relevant quantity is the deviation from this uniform baseline.

##### 1.2.1 Uniform line tension and effective preferred perimeter

For completeness, we show here how a uniform shift in line tension can be absorbed into the preferred perimeter, so that only *relative* line tensions matter for the mechanics. This justifies the choice $\gamma^{(c)}_{\text{hom}} = \gamma^{(b)}_{\text{hom}} = 0$ in the main text.

We start from the vertex energy

$$E = \sum_i \left[K_A(A_i - A_0)^2 + K_P(P_i - P_0)^2\right] + \sum_{\langle i,j\rangle} \gamma_{ij}\, l_{ij}, \tag{14}$$

where $A_i$ and $P_i$ are the area and perimeter of cell $i$, and $l_{ij}$ is the length of the edge shared by cells $i$ and $j$. The last term accounts for line tension along each cell–cell contact.

We now decompose the line tension into a spatially uniform baseline and a deviation,

$$\gamma_{ij} = \bar{\gamma} + \delta\gamma_{ij}, \tag{15}$$

with $\bar{\gamma}$ a constant and $\delta\gamma_{ij}$ capturing heterotypic and other edge-specific contributions. Inserting Eq. (15) into Eq. (14) gives

$$E = \sum_i \left[ K_A(A_i - A_0)^2 + K_P(P_i - P_0)^2 \right] + \bar{\gamma} \sum_{\langle i,j \rangle} l_{ij} + \sum_{\langle i,j \rangle} \delta\gamma_{ij}\, l_{ij}. \tag{16}$$

In a confluent tiling with periodic boundaries, each edge is shared by exactly two cells, so that the total edge length can be written as

$$\sum_{\langle i,j \rangle} l_{ij} = \frac{1}{2} \sum_i P_i. \tag{17}$$

Using Eq. (17), the baseline contribution becomes

$$\bar{\gamma} \sum_{\langle i,j \rangle} l_{ij} = \frac{\bar{\gamma}}{2} \sum_i P_i. \tag{18}$$

Collecting all perimeter-dependent terms for a given cell $i$, we obtain

$$K_P(P_i - P_0)^2 + \frac{\bar{\gamma}}{2} P_i = K_P P_i^2 - 2K_P P_0 P_i + K_P P_0^2 + \frac{\bar{\gamma}}{2} P_i. \tag{19}$$

This expression can be rewritten by completing the square,

$$K_P(P_i - P_0^{\text{eff}})^2 + \text{const}, \tag{20}$$

where the effective preferred perimeter is

$$P_0^{\text{eff}} = P_0 - \frac{\bar{\gamma}}{4K_P}. \tag{21}$$

The constant term does not affect forces or dynamics and can therefore be discarded. Thus a spatially uniform line tension $\bar{\gamma}$ simply renormalizes the preferred perimeter and shape index,

$$s_0^{\text{eff}} = \frac{P_0^{\text{eff}}}{\sqrt{A_0}} = s_0 - \frac{\bar{\gamma}}{4K_P\sqrt{A_0}}. \tag{22}$$

This argument shows that only deviations $\delta\gamma_{ij}$ from the uniform baseline enter the mechanical response. In particular, any common offset added to all $\gamma_{ij}$ leaves the dynamics unchanged up to a redefinition of $P_0$ and $s_0$. It is therefore sufficient to focus on relative line tensions, for instance by choosing a convention where all homotypic edges have zero tension and only heterotypic edges carry a nonzero contribution,

$$\gamma_{ij} = \begin{cases} \gamma_{\text{het}}, & \sigma_i \neq \sigma_j, \\ 0, & \sigma_i = \sigma_j. \end{cases} \tag{23}$$

as done in the main text. Positive $\gamma_{\text{het}}$ then penalizes cluster–background interfaces and favors their contraction, whereas negative $\gamma_{\text{het}}$ favors the expansion of heterotypic interfaces and promotes mixing between cluster and background, without any loss of generality from neglecting a global adhesion baseline.

**Simulation parameters and scan ranges.**

| Parameter | Meaning | Value / range | Used in |
| --- | --- | --- | --- |
| *Fixed model units* | | | |
| $A_0$ | preferred cell area | 1 | all simulations |
| $K_P$ | perimeter stiffness | 1 | all simulations |
| $K_A$ | area stiffness | 10 | all simulations |
| $\mu$ | mobility | 1 | all simulations |
| $L_x \times L_y$ | domain size | $20 \times 20$ | all simulations |
| $\Delta t$ | integration timestep | $2 \times 10^{-4}$ | all simulations |
| $t_{\text{eq}}$ | equilibration time | 1 | all simulations |
| $T$ | measurement window | 100 | all simulations |
| $\tau$ | polarity persistence time | 50 | all motility simulations |
| $D_r$ | rotational noise strength | 0.005 | all motility simulations |
| $n_{\text{sim}}$ | realizations per parameter set | 100 | all parameter scans |
| *Scanned parameters* | | | |
| $s_0$ | target shape index | [3.6, 4.0]; fixed to 3.7 in $\gamma$-scans | Main Figs. 1–2; Supp. Fig. S1 |
| $v_0$ | motility magnitude | [0.0, 4.0] | Main Figs. 1–3; Supp. Figs. S1–S5 |
| $\gamma_{\text{het}}$ | heterotypic line tension | [−1.5, 5.0] | Main Fig. 2; Supp. Figs. S2–S4 |
| $N_c$ | initial cluster size | $\{1, \dots, 8\}$ | Main Fig. 2; Supp. Figs. S2–S4 |
| $\alpha$ | adhesion scale | [0.0, 3.8] | Main Figs. 4–5; Supp. Figs. S8–S12 |
| $\beta$ | motility scale | [3.0, 8.8] | Main Figs. 4–5; Supp. Figs. S8–S12 |
| *AVM-specific* | | | |
| $\ell_{\min}$ | T1 edge threshold | 0.04 | AVM simulations |

### 2 Supplementary Note 2: Zebrafish-based parameterization of host tissue state and Nodal-dependent interactions

Zebrafish mesendoderm transplantation assays provide a setting in which the motility of donor cells and the mechanical state of the host tissue can be varied independently during gastrulation [5]. Here we use these experiments to parameterize the vertex model in two steps: first, by estimating the geometric state of the ectodermal host tissue from membrane segmentation; and second, by mapping experimentally measured Nodal signalling levels to cell motility and interfacial line tensions.

#### 2.1 Ectoderm shape measurements and inference of host tissue state

The image-processing steps used to obtain ectoderm cell outlines are described in the Methods. Here, we focus on how those measurements constrain the background tissue state used in the model. Because the vertex model frameworks used here are two-dimensional, we quantified cell geometry from single optical sections selected from z-stacks, yielding polygonal cell outlines that can be compared directly with the vertex description.

A limitation of this approach is that the tissue itself is three-dimensional, so two-dimensional optical sections inevitably introduce sectioning effects. However, Sharp et al. demonstrated that 2D cell-shape distributions measured in slices through a 3D tissue retain informative statistical signatures of the underlying 3D shape state, making 2D shape statistics a meaningful proxy for tissue geometry [6].

In particular, following this work, we denote by $s \equiv P_{\text{measured}}/\sqrt{A_{\text{measured}}}$ the measured projected cell-shape index extracted from segmented cell outlines. Because optical-sectioning effects broaden the measured distributions and generate a high-$s$ tail [6], especially for cells that are only partially intersected by the imaging plane, we summarize the data using the peak of the $s$ distribution rather than the mean or median. Across the analysed time window, this peak remained close to $s_{\text{peak}} \approx 3.81$ (Fig. 4d; Supp. Fig. S6), with a small increase over time.

In the SPV framework, the rigidity transition is controlled by the preferred shape index $s_0$ and occurs near $s_0^* \approx 3.81$ [3]. Importantly, the measured shape index $s$ is not identical to the input parameter $s_0$: In the solid regime ($s_0 < s_0^*$), the typical observed cell shape $s$ remains close to the critical value, whereas in the fluid regime ($s_0 > s_0^*$), the observed shape increases further and more closely tracks the input $s_0$ [3, 7]. We therefore use the measured peak shape index as a geometrically motivated indicator of host-tissue state. A measured value close to, but not clearly above, $s \approx 3.81$ is thus consistent with a host tissue that lies in a jammed, solid-like regime [8].

In the simulations, we therefore chose $s_0 = 3.7$ as a conservative background value. This places the host tissue on the solid-like side of the transition while remaining close enough to the threshold that sufficiently strong active forces can still induce local rearrangements and invasion, consistent with the zebrafish transplantation setting in Ref. [5].

#### 2.2 Mapping Nodal signalling to motility and line tension

We next map Nodal signalling to the motility and interfacial parameters of the vertex model. As in the active-particle framework of Pinheiro et al. [5], each cell is assigned a scalar Nodal level $N$. In the vertex model, this signalling level sets both the active motility magnitude $v_0(N)$ and the line tension $\gamma_{ij}(N_i, N_j)$ along interfaces between cells $i$ and $j$. Following previous minimal mechano-chemical descriptions of tissues [9–11], we seek the simplest mapping that captures the transplantation hierarchy.

##### 2.2.1 Experimental Nodal levels used for donor populations

Effective Nodal levels were obtained by re-analysing the nuclear pSmad2/3 intensity profiles reported in Ref. [5] for donor populations taken from 50% epiboly (Nodal High), shield stage (Nodal Medium), and 75% epiboly (Nodal Low) embryos. To match the region from which donor cells are collected in the transplantation assays, we restricted the analysis to the first three cell tiers from the dorsal margin (tiers 0–2; Supp. Fig. 8a). This yields the effective signalling levels

$$N_{\text{High}} \approx 0.44, \qquad N_{\text{Medium}} \approx 0.20, \qquad N_{\text{Low}} \approx 0.08.$$

In the simulations, donor cells are therefore assigned one of these three discrete Nodal levels, while the MZoep host tissue is assigned $N_{\text{host}} = 0$.

##### 2.2.2 Motility rule: $v_0 = \beta N$

Pinheiro et al. [5] showed that Nodal signalling correlates with protrusive activity during mesendoderm internalization. We therefore assign each cell $i$ an active motility magnitude

$$v_{0,i} = \beta N_i,$$

where $\beta$ is a global motility scale inferred from the transplant data. Under this rule, 50% epiboly donor cells are the most motile, shield-stage donor cells are less motile, 75% epiboly donor cells are weaker still, and host ectodermal cells with $N_{\text{host}} = 0$ are passive.

##### 2.2.3 Adhesion rules

To capture the Nodal dependence of cell–cell interactions, we compare two minimal line-tension rules. In both cases, the experimentally inferred Nodal levels are used directly, so that once $(\alpha, \beta)$ are fixed, all donor–donor and donor–host interfaces are determined.

As a vertex analogue of a heterotypic interaction or preferential adhesion rule, we assign

$$\gamma_{ij}^{\text{int}} = \alpha|N_i - N_j|,$$

with $\alpha > 0$. Under this rule, homotypic contacts carry no additional line tension, while heterotypic contacts carry positive line tension that increases with the Nodal difference between the two cells. Interfaces between cells with similar Nodal levels are therefore effectively more cohesive than interfaces between cells with strongly different Nodal levels. In the vertex/SPV setting, this is the natural extension of the heterotypic cluster–background line tension considered in the main text.

As a vertex analogue of a differential adhesion rule, we instead assign

$$\gamma_{ij}^{\text{hom}} = -\alpha\sqrt{N_i N_j},$$

again with $\alpha > 0$. In this case, interfaces between highly signalling cells acquire more negative line tension, corresponding to stronger effective adhesion, whereas interfaces involving weakly signalling or host cells are less cohesive. This construction mirrors differential-adhesion formulations in particle and vertex models [12–14], while remaining compatible with the effective edge-energy description used in SPV and AVM [4, 15, 16].

Taken together, the two relations

$$\gamma_{ij}^{\text{int}} = \alpha|N_i - N_j| \qquad \text{and} \qquad \gamma_{ij}^{\text{hom}} = -\alpha\sqrt{N_i N_j}$$

define the two Nodal-based interaction rules tested in the model.

### 3 Supplementary Note 3: Constraining model parameters with zebrafish transplant assays

Zebrafish transplantation assays provide a hierarchy of experimental constraints on the two free parameters of the model, the motility scale $\beta$ and the interaction scale $\alpha$. Rather than fitting a single condition in isolation, we required a candidate parameter pair $(\alpha, \beta)$ to reproduce the ordering of outcomes across homogeneous, mixed, and triple donor populations. The corresponding simulation observables and agreement criteria are defined in the Methods; here we focus on how the successive transplant classes restrict parameter space and distinguish between the two Nodal-dependent interaction rules.

#### 3.1 Median-based definition of best-fit regions

Near the boundaries between migratory and non-migratory regimes, simulation outcomes are often broad and skewed across realizations, so mean values can be strongly influenced by a minority of runs with unusually strong invasion or unusually extensive breakup. Supp. Fig. 11

illustrates this clearly for several mixed-cluster observables, where mean and median summaries differ substantially over the same parameter range. A similar logic applies to the transplantation assays of Pinheiro et al. [5], whose outcomes are themselves heterogeneous across embryos and donor clusters. We therefore defined best-fit regions from the median rather than the mean, so that the selected parameter regions capture the typical outcome rather than being biased by rare events. This is particularly important for mixed and triple transplants, where stochastic fragmentation and partial co-migration can coexist near the transition between coordinated and uncoordinated behaviour.

#### 3.2 Hierarchical constraints from homogeneous and mixed transplants

Homogeneous donor populations provide the first coarse constraint on the parameter plane. In practice, these assays define an admissible region in $(\alpha, \beta)$ space rather than constraining only a single parameter. The requirement that High donors internalize while Medium and Low donors remain below the internalization threshold imposes a strong restriction on the motility scale $\beta$. At the same time, the donor–host interface is governed by the same non-monotonic invasion logic identified by the model: when $\alpha$ is too small, donor clusters are insufficiently cohesive and split, whereas when $\alpha$ is too large, donor–host coupling suppresses efficient advance. The homogeneous transplants therefore already select a broad intermediate band of $\alpha$ values, as seen in Supp. Fig. 9.

Mixed transplants then provide a more stringent test of coupling between leader and follower populations. Experimentally, Nodal High+Medium clusters remain cohesive and co-migrate, whereas Nodal High+Low clusters frequently lose their Nodal Low followers [5]. These outcomes sharpen the admissible parameter window considerably. If $\alpha$ is too small, both mixed clusters tend to fragment, with only the most motile donors advancing. If $\alpha$ is too large, the less motile followers oppose leader advance strongly enough to suppress collective internalization. Parameter values that reproduce both the High+Medium and High+Low behaviours therefore lie in a narrower intermediate window, obtained by intersecting the mixed-cluster constraints with the broader region already selected by the homogeneous assays (Fig. 4; Supp. Fig. 10).

#### 3.3 Triple transplants distinguish the interaction rules

Triple transplants containing (Nodal-) High, Medium, and Low donor populations provide the most rigorous test of the interaction rules, because they require a single parameter pair $(\alpha, \beta)$ to reproduce not only invasion, but also coupling across the full Nodal hierarchy. Experimentally, inserting Medium cells between High and Low populations partially rescues collective behaviour, allowing a substantial fraction of low-Nodal cells to internalize together with the cluster [5]. The relevant model requirement is therefore not simply that triple clusters move, but that they do so while retaining sufficient cohesion across all three donor populations.

Under the heterotypic rule,

$$\gamma_{ij}^{\text{int}} = \alpha |N_i - N_j|,$$

this behaviour emerges naturally. Parameter values that already satisfy the homogeneous and mixed-cluster best-fit regions continue to admit triple-population internalization with limited breakup (Fig. 5; Supp. Fig. 12a–c). In this regime, the experimentally observed rescue is reproduced because contacts between nearby Nodal levels remain sufficiently cohesive to maintain cluster-wide coupling. In particular, High–Medium and Medium–Low interfaces both fall into an intermediate coupling regime, so Medium cells can transmit motion from highly motile High leaders to weakly motile Low followers without creating a destabilizing weak link within the cluster.

Under the differential adhesion rule,

$$\gamma_{ij}^{\text{hom}} = -\alpha \sqrt{N_i N_j},$$

the same rescue is much harder to obtain. Although parts of the homogeneous and mixed-cluster hierarchy can still be reproduced, the overlap region that also satisfies the triple-transplant constraints disappears altogether (Fig. 5; Supp. Figs. 9g–l, 10g–l, and 12d–f). In this case, no single parameter pair robustly supports both the required leader–follower cohesion and the partial rescue of low-Nodal cells in triple clusters.

The distinction between the two rules follows directly from how they distribute cohesion across the Nodal hierarchy. Under the heterotypic interaction rule, line tension is set by differences in Nodal level, so mechanically similar neighbours remain coupled: High stays linked to Medium, Medium stays linked to Low, and the cluster can therefore internalize as a connected chain even though High and Low differ markedly from one another. Under the differential adhesion rule, by contrast, cohesion is concentrated among the cells with higher Nodal levels. High–Medium contacts are then favored over Medium–Low contacts, making the Medium–Low interface the weakest donor–donor link in the cluster. As a result, Medium preferentially remains associated with High, whereas Low is more easily left behind when the cluster advances. In this sense, the failure of the differential adhesion rule is not simply that adhesion is globally too weak or too strong, but that it places the weakest link exactly where mechanical continuity is required for rescue. The resulting difference in interaction logic is summarized schematically in Fig. 5i and explains why the heterotypic interaction rule, but not the differential adhesion one, robustly reproduces the triple-transplant behaviour.

### Appendices

#### A SPV force derivations

This section collects the explicit geometric derivatives used to evaluate the SPV forces in Eqs. (7)–(8). The notation follows the construction in Extended Fig. 1: each Voronoi vertex $\mathbf{h}_{ijk}$ is the circumcenter of a Delaunay triangle $(i, j, k)$ where three cells meet.

For a given cell $c \in \{i, j, k\}$, denote its consecutive Voronoi vertices along the boundary by

$$\boldsymbol{\nu}^{(c)}_{\text{prev}}, \quad \boldsymbol{\nu}^{(c)}_{0} = \mathbf{h}_{ijk}, \quad \boldsymbol{\nu}^{(c)}_{\text{next}}.$$

Define the local edge vectors and their unit tangents as

$$\mathbf{e}^{(c)}_{\text{prev}} = \boldsymbol{\nu}^{(c)}_{0} - \boldsymbol{\nu}^{(c)}_{\text{prev}}, \qquad \mathbf{e}^{(c)}_{\text{next}} = \boldsymbol{\nu}^{(c)}_{\text{next}} - \boldsymbol{\nu}^{(c)}_{0},$$

$$\hat{\mathbf{t}}^{(c)}_{\text{prev}} = \frac{\mathbf{e}^{(c)}_{\text{prev}}}{\|\mathbf{e}^{(c)}_{\text{prev}}\|}, \qquad \hat{\mathbf{t}}^{(c)}_{\text{next}} = \frac{\mathbf{e}^{(c)}_{\text{next}}}{\|\mathbf{e}^{(c)}_{\text{next}}\|}.$$

Using the rotation operator $\mathsf{J}(x, y) = (-y, x)$, standard polygon calculus gives the sensitivities of area and perimeter to the vertex position:

$$\frac{\partial A_c}{\partial \mathbf{h}_{ijk}} = \tfrac{1}{2}\,\mathsf{J}\Big(\boldsymbol{\nu}^{(c)}_{\text{next}} - \boldsymbol{\nu}^{(c)}_{\text{prev}}\Big), \tag{24}$$

$$\frac{\partial P_c}{\partial \mathbf{h}_{ijk}} = \hat{\mathbf{t}}^{(c)}_{\text{prev}} - \hat{\mathbf{t}}^{(c)}_{\text{next}}. \tag{25}$$

From these relations, the cell-level contribution to the bulk energy derivative is

$$\frac{\partial E_c}{\partial \mathbf{h}_{ijk}} = 2K_A(A_c - A_0)\,\frac{\partial A_c}{\partial \mathbf{h}_{ijk}} + 2K_P(P_c - P_0)\,\frac{\partial P_c}{\partial \mathbf{h}_{ijk}}. \tag{26}$$

Summing over the three cells that meet at the vertex gives the bulk contribution entering Eq. (7).

For the interfacial term, the three edges $(i, j)$, $(j, k)$, and $(k, i)$ that converge at $\mathbf{h}_{ijk}$ contribute $\gamma_{ab}\ell_{ab}$ each. Let $\hat{\mathbf{t}}^{(ab)}$ be the unit tangent of edge $(a, b)$ oriented away from the vertex. Since

$$\frac{\partial \ell_{ab}}{\partial \mathbf{h}_{ijk}} = -\hat{\mathbf{t}}^{(ab)},$$

the derivative of the interfacial energy reads

$$\frac{\partial E_{\text{tens}}}{\partial \mathbf{h}_{ijk}} = -\,\gamma_{ij}\,\hat{\mathbf{t}}^{(ij)} - \gamma_{jk}\,\hat{\mathbf{t}}^{(jk)} - \gamma_{ki}\,\hat{\mathbf{t}}^{(ki)}. \tag{27}$$

Inserted into Eq. (6), this yields the interfacial force contribution in Eq. (8).

The sensitivity of a Voronoi vertex to the displacement of a cell center follows from the circumcenter of the Delaunay triangle. For $(i, j, k)$, define

$$\mathbf{r}_{ij} = \mathbf{r}_j - \mathbf{r}_i, \qquad \mathbf{r}_{ik} = \mathbf{r}_k - \mathbf{r}_i, \qquad \mathbf{r}_{jk} = \mathbf{r}_k - \mathbf{r}_j,$$

and the auxiliary quantities

$$c = r_{ij,x}\, r_{jk,y} - r_{ij,y}\, r_{jk,x}, \qquad d = 2c^2, \tag{28}$$

$$\beta d = -\|\mathbf{r}_{ik}\|^2 (\mathbf{r}_{ij}\cdot\mathbf{r}_{jk}), \qquad \gamma d = \|\mathbf{r}_{ij}\|^2 (\mathbf{r}_{ik}\cdot\mathbf{r}_{jk}), \tag{29}$$

$$\mathbf{z} = \beta d\, \mathbf{r}_{ij} + \gamma d\, \mathbf{r}_{ik}. \tag{30}$$

The Jacobian of the circumcenter with respect to the cell-center position is

$$\frac{\partial \mathbf{h}_{ijk}}{\partial \mathbf{r}_i} = I_2 + \frac{1}{d}\Big[\mathbf{r}_{ij}\otimes\partial_{\mathbf{r}_i}(\beta d) + \mathbf{r}_{ik}\otimes\partial_{\mathbf{r}_i}(\gamma d) - (\beta d + \gamma d) I_2 - \mathbf{z}\otimes\Big(\frac{1}{d}\,\partial_{\mathbf{r}_i} d\Big)\Big], \tag{31}$$

where

$$\partial_{\mathbf{r}_i}(\beta d) = 2(\mathbf{r}_{ij}\cdot\mathbf{r}_{jk})\,\mathbf{r}_{ik} + \|\mathbf{r}_{ik}\|^2\,\mathbf{r}_{jk}, \tag{32}$$

$$\partial_{\mathbf{r}_i}(\gamma d) = -2(\mathbf{r}_{ik}\cdot\mathbf{r}_{jk})\,\mathbf{r}_{ij} + \|\mathbf{r}_{ij}\|^2\,\mathbf{r}_{jk}, \tag{33}$$

$$\frac{1}{d}\,\partial_{\mathbf{r}_i} d = \frac{2}{c}\,(-r_{jk,y},\, r_{jk,x}). \tag{34}$$

When the three points become nearly colinear ($c \to 0$), the circumcenter is ill-conditioned, but in simulations the Delaunay triangulation is recomputed at each step, and edge flips resolve such degeneracies automatically. Therefore, Eq. (31) is evaluated only away from the singular limit.

Substituting Eqs. (24)–(26), Eq. (27), and Eq. (31) into Eq. (6) yields the compact force expressions in Eqs. (7) and (8) used for the SPV dynamics.

### B AVM force derivations

This section collects the explicit geometric derivatives used to evaluate the AVM forces in Eqs. (9)–(10). Each vertex $\boldsymbol{\nu}$ is shared by up to three neighboring cells $(i, j, k)$, as illustrated in Extended Fig. 2. All quantities are expressed in terms of the local vertex configuration. Vertices are the independent degrees of freedom, so the mechanical force on each vertex is

$$\mathbf{F}(\boldsymbol{\nu}) = -\frac{\partial E}{\partial \boldsymbol{\nu}} = \mathbf{F}_{\text{bulk}}(\boldsymbol{\nu}) + \mathbf{F}_{\text{tens}}(\boldsymbol{\nu}),$$

without the circumcenter Jacobian present in SPV.

For a given cell $i$, let the consecutive vertices along its boundary be

$$\boldsymbol{\nu}^{(i)}_{\text{prev}}, \quad \boldsymbol{\nu}^{(i)}_{0} = \boldsymbol{\nu}, \quad \boldsymbol{\nu}^{(i)}_{\text{next}},$$

and define the corresponding edge vectors and unit tangents

$$\mathbf{e}^{(i)}_{\text{prev}} = \boldsymbol{\nu} - \boldsymbol{\nu}^{(i)}_{\text{prev}}, \qquad \mathbf{e}^{(i)}_{\text{next}} = \boldsymbol{\nu}^{(i)}_{\text{next}} - \boldsymbol{\nu},$$

$$\hat{\mathbf{t}}^{(i)}_{\text{prev}} = \frac{\mathbf{e}^{(i)}_{\text{prev}}}{\|\mathbf{e}^{(i)}_{\text{prev}}\|}, \qquad \hat{\mathbf{t}}^{(i)}_{\text{next}} = \frac{\mathbf{e}^{(i)}_{\text{next}}}{\|\mathbf{e}^{(i)}_{\text{next}}\|}.$$

Using the rotation operator $\mathsf{J}(x, y) = (-y, x)$, the sensitivities of area and perimeter to the vertex position are

$$\frac{\partial A_i}{\partial \boldsymbol{\nu}} = \tfrac{1}{2}\,\mathsf{J}\big(\boldsymbol{\nu}^{(i)}_{\text{next}} - \boldsymbol{\nu}^{(i)}_{\text{prev}}\big), \tag{35}$$

$$\frac{\partial P_i}{\partial \boldsymbol{\nu}} = \hat{\mathbf{t}}^{(i)}_{\text{prev}} - \hat{\mathbf{t}}^{(i)}_{\text{next}}, \tag{36}$$

which define how the area and perimeter of cell $i$ respond to infinitesimal vertex displacements.

The contribution of cell $i$ to the bulk energy gradient is

$$\frac{\partial E_i}{\partial \boldsymbol{\nu}} = 2K_A(A_i - A_0)\,\frac{\partial A_i}{\partial \boldsymbol{\nu}} + 2K_P(P_i - P_0)\,\frac{\partial P_i}{\partial \boldsymbol{\nu}}. \tag{37}$$

Summing over the three cells that share the vertex gives

$$\frac{\partial E_{\text{bulk}}}{\partial \boldsymbol{\nu}} = \frac{\partial E_i}{\partial \boldsymbol{\nu}} + \frac{\partial E_j}{\partial \boldsymbol{\nu}} + \frac{\partial E_k}{\partial \boldsymbol{\nu}}. \tag{38}$$

Applying $\mathbf{F}_{\text{bulk}}(\boldsymbol{\nu}) = -\,\partial E_{\text{bulk}}/\partial \boldsymbol{\nu}$ then gives Eq. (9).

For the interfacial term, the edges meeting at $\boldsymbol{\nu}$ correspond to interfaces $(i, j)$, $(j, k)$, and $(k, i)$. By convention, the edge along $\hat{\mathbf{t}}^{(i)}_{\text{next}}$ separates cells $i$ and $j$, and the edge along $\hat{\mathbf{t}}^{(i)}_{\text{prev}}$ separates cells $i$ and $k$. The derivatives of the corresponding edge lengths with respect to the vertex position are

$$\frac{\partial \ell_{ij}}{\partial \boldsymbol{\nu}} = -\,\hat{\mathbf{t}}^{(i)}_{\text{next}}, \qquad \frac{\partial \ell_{ik}}{\partial \boldsymbol{\nu}} = +\,\hat{\mathbf{t}}^{(i)}_{\text{prev}}.$$

The contribution of cell $i$ to the interfacial energy derivative is then

$$\left.\frac{\partial E_{\text{tens}}}{\partial \boldsymbol{\nu}}\right|_i = \gamma_{ik}\,\hat{\mathbf{t}}^{(i)}_{\text{prev}} - \gamma_{ij}\,\hat{\mathbf{t}}^{(i)}_{\text{next}}. \tag{39}$$

Adding the cyclic permutations $i \to j \to k \to i$ gives

$$\frac{\partial E_{\text{tens}}}{\partial \boldsymbol{\nu}} = (\gamma_{ik}\,\hat{\mathbf{t}}^{(i)}_{\text{prev}} - \gamma_{ij}\,\hat{\mathbf{t}}^{(i)}_{\text{next}}) + (\gamma_{ji}\,\hat{\mathbf{t}}^{(j)}_{\text{prev}} - \gamma_{jk}\,\hat{\mathbf{t}}^{(j)}_{\text{next}}) + (\gamma_{kj}\,\hat{\mathbf{t}}^{(k)}_{\text{prev}} - \gamma_{ki}\,\hat{\mathbf{t}}^{(k)}_{\text{next}}), \tag{40}$$

and therefore, after applying $\mathbf{F}_{\text{tens}}(\boldsymbol{\nu}) = -\,\partial E_{\text{tens}}/\partial \boldsymbol{\nu}$, yields Eq. (10).

Combining these results gives the total vertex force,

$$\mathbf{F}(\boldsymbol{\nu}) = \mathbf{F}_{\text{bulk}}(\boldsymbol{\nu}) + \mathbf{F}_{\text{tens}}(\boldsymbol{\nu}), \tag{41}$$

$$\mathbf{F}_{\text{bulk}}(\boldsymbol{\nu}) = -\left(\frac{\partial E_i}{\partial \boldsymbol{\nu}} + \frac{\partial E_j}{\partial \boldsymbol{\nu}} + \frac{\partial E_k}{\partial \boldsymbol{\nu}}\right), \tag{42}$$

$$\mathbf{F}_{\text{tens}}(\boldsymbol{\nu}) = -\,\frac{\partial E_{\text{tens}}}{\partial \boldsymbol{\nu}}, \tag{43}$$

with $\partial E_{\text{tens}}/\partial \boldsymbol{\nu}$ given by Eq. (40).

The vertex dynamics follow directly from the expressions above. Substituting Eqs. (35)–(37) and Eq. (40) into Eqs. (42) and (43) yields the compact force forms in Eqs. (9) and (10). The vertex positions are then updated according to Eq. (3), with $\mathbf{x}_i = \boldsymbol{\nu}$ and the active term $\mathbf{v}_i = \mathbf{v}_{\text{act}}(\boldsymbol{\nu})$ taken as the average of the propulsive contributions of the cells meeting at that vertex. All geometric derivatives are evaluated directly from the instantaneous polygon geometry at every simulation step.
